# Spatiotemporal profiling reveals emergence of cell type-specific molecular aging and senescence programs in the spinal cord

**DOI:** 10.64898/2026.11.17.767384

**Authors:** Olena Kuksenko, Maya Xia, Aidan C. Daly, Christopher A. Jackson, Natalie Barretto, Imdadul Haq, Mark A. Watson, Christina D. King, Mariela Bejan, Patricia Santos, Courteney Mattison, Maria Hauge Pedersen, Neil A. Shneider, Vilas Menon, Birgit Schilling, Hemali Phatnani

## Abstract

The spinal cord is the primary relay center between the periphery and the brain, playing a critical role in sensory, motor and autonomic function. Despite its functional importance, the cellular and molecular changes underlying spinal cord aging and their relationship to cellular senescence – a hallmark associated with aging – remain incompletely understood. Here, we generated a cross-species, multi-omic atlas of spinal cord aging in mice and humans by integrating a combination of liquid chromatography-tandem mass spectrometry, Visium spatial transcriptomics, single-nucleus RNA-sequencing, and multiplexed immunofluorescence imaging. We identified converging age-associated increases in fibrinogen, laminin, and immune activation-associated proteins across the mouse and human spinal cord. Over the mouse lifespan, we further identified distinct spatiotemporal patterns of aging. Downregulated transcriptional programs, including synaptic signaling, mitochondrion organization, and autophagy, were most highly expressed in grey matter regions and in neurons. Upregulated transcriptional programs, including vascular and immune processes, were enriched in white matter regions and in endothelial cells and microglia, respectively. Notably, these increasing spatial modules were also enriched for senescence-associated genes. In the human spinal cord, endothelial cells and microglia were also most enriched for canonical senescence markers *CDKN1A* and *CDKN2A*, respectively. Overall, this study defines the shared, spatially-resolved, cell type-specific features associated with aging and senescence in the mouse and human spinal cord.

## Introduction

The spinal cord relays information between the brain and the periphery, coordinating essential functions including movement, respiration, autonomic regulation, and sensation. Extending caudally from the brainstem through the length of the vertebral column, the spinal cord has a highly organized architecture that differs significantly from the brain, where grey matter surrounds the interior white matter^1^. In contrast, the spinal cord’s external white matter is composed of myelinated axonal tracts carrying ascending sensory and descending motor information, while the internal grey matter contains motor neurons in the ventral horn and sensory processing interneurons in the dorsal horn. The spinal cord is susceptible to aging^2–6^ and to age-associated neurodegenerative diseases, including amyotrophic lateral sclerosis^7–10^. Despite its functional importance, the cellular and molecular changes that underlie aging in the spinal cord remain poorly understood.

One major hallmark of aging is cellular senescence^11–13^, which is a cellular state triggered by prolonged cell stress and characterized by stable, essentially irreversible cell cycle arrest^12,14^. Senescence is primarily enforced by two tumor-suppressive pathways, p53-p21 (*CDKN1A*) and p16 (*CDKN2A*)-pRB, which converge downstream on the inhibition of CDK2 and CDK4/6, together blocking progression from G1 to S phase^15–19^. Senescent cells also release senescence-associated secretory phenotype (SASP) factors that alter the tissue microenvironment and contribute to aging^20–22^. SASP includes pro-inflammatory cytokines, chemokines, exosomes, growth factors, and extracellular matrix-degrading proteins. Broadly, senescent cells have been implicated in physiological functions including embryonic development, inflammation, wound healing, and tumor suppression^15,23–25^. A growing body of literature suggests that senescent cells also increase with aging and age-associated diseases, including in the central nervous system, with p16 and p21 used as proxy markers to identify these cells^12,14,26–31^. Moreover, clearing these cells using pharmacological tools such as senolytics ameliorates age-associated phenotypes in mice^28,29,32,33^. Senescent cells have also been implicated in mouse models of spinal cord injury^34^, chronic pain^35^, and amyotrophic lateral sclerosis^36–39^, and senolytic treatment improves recovery from spinal cord injury^40^, underscoring the importance of understanding these cells in the spinal cord.

There are many challenges associated with characterizing senescent cells since their markers are context- and cell type-dependent, and no universal markers exist, requiring unbiased approaches to profile these cells^31,41–45^. Using spatial transcriptomics, single nucleus RNA sequencing, and multiplexed protein imaging, our lab previously identified cell type-specific aging and senescence signatures in the human dorsolateral prefrontal cortex (DLPFC) and hippocampus across two aging cohorts of non-neuropathological donors^46,47^. In the DLPFC, we found that the white matter is a hotspot for senescence-associated signatures, and p16/*CDKN2A* was enriched in oligodendrocytes and increased with age^46^. In the hippocampus, senescence-associated genes were enriched in microglial and neurovascular programs, and p16/*CDKN2A* was enriched in oligodendrocytes while p21/*CDKN1A* was enriched in endothelial cells^47^. Across both regions, p16 and p21 cells were non-overlapping cell populations. However, given the differences in cell type composition and architecture between the spinal cord and the brain, where and in which cell types senescent signatures arise in the spinal cord remain poorly defined.

In this study, we mapped signatures of aging and senescence in the mouse and human spinal cord using liquid chromatography-tandem mass spectrometry (LC-MS/MS), spatial transcriptomics (ST), single nucleus RNA sequencing (snRNA-seq), and multiplexed protein imaging. Across species, we found convergent increases in age-associated signatures, including in fibrinogen, extracellular matrix, and immune activation proteins. In mice, aging-related changes included a downregulation (e.g. synaptic signaling, mitochondrion organization, autophagy) and upregulation (e.g. circulatory system, immune system processes) in several spatial gene modules. Interestingly, the circulatory system and immune system modules were primarily expressed in the white matter and enriched for senescence-associated genes. Finally, in the human spinal cord, p21/*CDKN1A* was enriched in endothelial cells and microglia, and p16/*CDKN2A* was enriched in microglia and oligodendrocytes. Together, these findings define shared, spatially resolved, cell type-specific features associated with aging and senescence in the spinal cord across species.

## Results

### A multi-omic atlas of the aging mouse and human spinal cord

To characterize the molecular signatures of aging and cellular senescence in the spinal cord, we employed an integrative multi-omics approach combining mass spectrometry-based proteomics (LC-MS/MS), spatial transcriptomics (ST), and multiplexed immunofluorescence (IF) in mice, with LC-MS/MS, single-nucleus RNA sequencing (snRNA-seq), and multiplexed immunofluorescence in human postmortem tissue (Figure 1A, Table S1). We performed mass spectrometry-based proteomic profiling on spinal cord tissue from 16 male mice spanning four age groups (6, 12, 15, and 24 months; n = 4 per age group). In parallel, we performed ST using the 10X Genomics Visium platform on lumbar spinal cord tissue from 20 mice across five age groups (6, 12, 18, 24, and 30 months), with four mice per age group (two males and two females), and four tissue sections per mouse.

**Figure 1.**
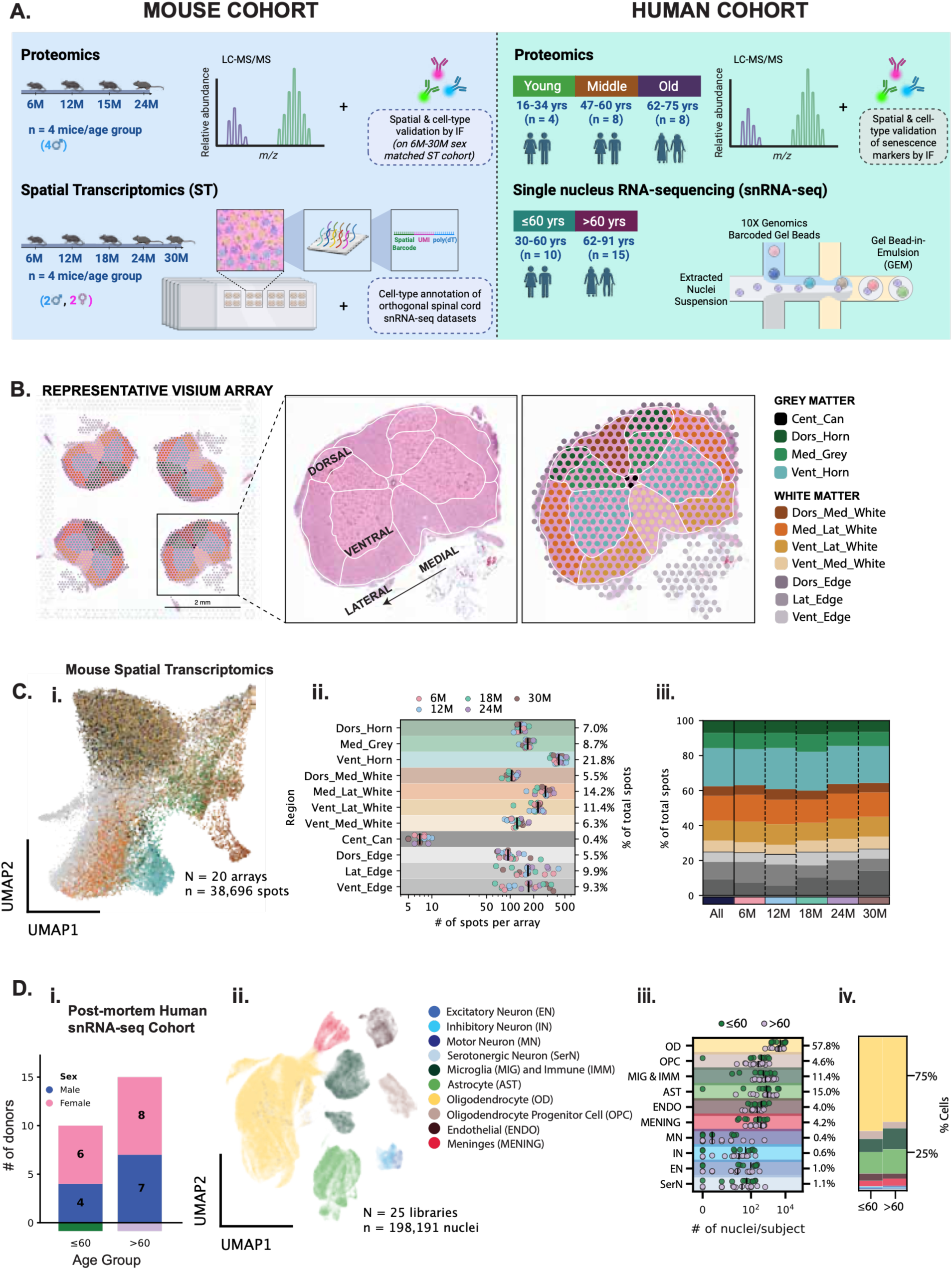
A multi-omic atlas of the aging mouse and human spinal cord. **A)** Overview of aging mouse and non-neuropathological human spinal cord cohorts for proteomics (LC-MS/MS, multiplexed immunofluorescence) and gene expression maps (ST, snRNA-seq). **B)** Representative image of an annotated mouse Visium spatial transcriptomics array (4 spinal cords/array). Each spot was assigned an annotated anatomical region (AAR) based on tissue morphology, cellular composition, and anatomical landmarks as seen in the H&E staining. **C)** i) Uniform manifold approximation and projection (UMAP) plot of all analyzed ST data (N=20 arrays, n=38,696 spots). Each dot represents a spot colored by AAR. ii) Number of spots annotated per AAR across Visium arrays, colored by age. iii) Relative proportion of spots across spinal cord AARs separated by age group. No significant differences were found in spot proportions. **D)** i) Human donor distribution of age groups split by sex. ii) UMAP of all analyzed snRNA-seq data (N=25 libraries, n=198,191 nuclei). Each dot represents a nucleus colored by its cell type. iii) Number of nuclei annotated per cell type per subject colored based on age group, after quality control and doublet exclusion. Percentage shows the proportion of nuclei assigned to each cell type. Black bar marks median nuclei counts. iv) Relative proportion of nuclei per cell type split by age group. No significant differences were found in cell type proportions.

Each Visium capture array contained 4,992 spatially barcoded RNA capture spots (55 µm in diameter), accommodating four spinal cord tissue sections per array. Across the 20 arrays comprising 80 tissue sections, we obtained transcriptomic profiles from 38,696 spatial spots, with a mean of 3641 unique RNA molecules and 1706 genes per spot (Figure S1, Table S2). To enable region-specific analyses, we classified each spatial transcriptomic spot into 11 anatomical annotation regions (‘AARs’) using hematoxylin and eosin (H&E) staining, cellular composition and anatomical landmarks as guides (Figure 1B)^48^. Visualization of preprocessed spatial transcriptomics spots using Uniform Manifold Approximation and Projection (UMAP) revealed a distinct separation of the central canal (Cent_Can) from the surrounding grey and white matter regions (Figure 1Ci). Each of the 11 AARs were consistently represented across all age groups, sexes and individual mice, and this regional composition was preserved with age (Figure 1Cii-iii, Figure S1). To enable the identification of cell-type contributions to spatially resolved transcriptional programs, we compiled five publicly available spinal cord snRNA-seq datasets^49–53^ with complementary cellular representation, including neuron-enriched datasets, and uniformly reprocessed them to generate a combined cell-type reference (Figure S2, Table S3). Finally, we validated our findings using multiplexed immunofluorescence across the same cohort of mice analyzed by spatial transcriptomics.

The post-mortem human cohort consisted of lumbar spinal cord tissue from sex-balanced, aging donors (Table S1). The LC-MS/MS cohort comprised 20 donors aged 16-75 years, the snRNA-seq cohort included 25 donors aged 30-91 years, and the IF cohort comprised 13 donors aged 45-78 years (Figure 1Di; Table S1). To investigate the cellular landscape of the aging human spinal cord, we performed cell-type annotation of the snRNA-seq dataset using gene expression profiles and canonical cell-type specific markers from the literature (Figure 1Dii, Table S3,4). Following the removal of doublets and low-quality nuclei, we assigned the remaining 198,191 nuclei into 10 major CNS cell types: excitatory neurons (EN), inhibitory neurons (IN), motor neurons (MN), serotonergic neurons (SerN), oligodendrocytes (OD), oligodendrocyte progenitor cells (OPC), astrocytes (AST), microglia and immune (MIG/IMM), endothelial (ENDO), meninges (MENING)^54^. The cellular composition of this dataset was dominated by glial cells, in particular oligodendrocytes (57.8%), astrocytes (15%), and microglia (11.4%) (Figure 1Diii). All major cell types were represented across age groups and sexes and were detected in most donors (Figure 1Diii, Figure S4). The relative proportions of major cell types did not change significantly with age (Figure 1Div, Figure S3). To complement our snRNA-seq analyses, we used multiplexed immunofluorescence to examine the expression and cellular localization of senescence-associated markers. Together, these combined proteomic and transcriptomic modalities provide a framework for defining aging and senescence signatures across species.

### Global age-associated proteomic changes in the mouse and human spinal cord

We first wanted to investigate the global changes associated with spinal cord aging. We performed quantitative liquid chromatography-tandem mass spectrometry (LC-MS/MS) on whole spinal cord tissue from male mice at 6, 12, 15, and 24 months of age (n = 4 per age group; Figure 2A). Following protein extraction and tryptic digestion, peptides were analyzed using data-independent acquisition (DIA) mass spectrometry, and protein abundances were quantified using Spectronaut. This approach enabled the quantification of 7,751 protein groups across the mouse spinal cord samples, providing a global overview of age-associated changes in the spinal cord proteome (Table S5).

**Figure 2.**
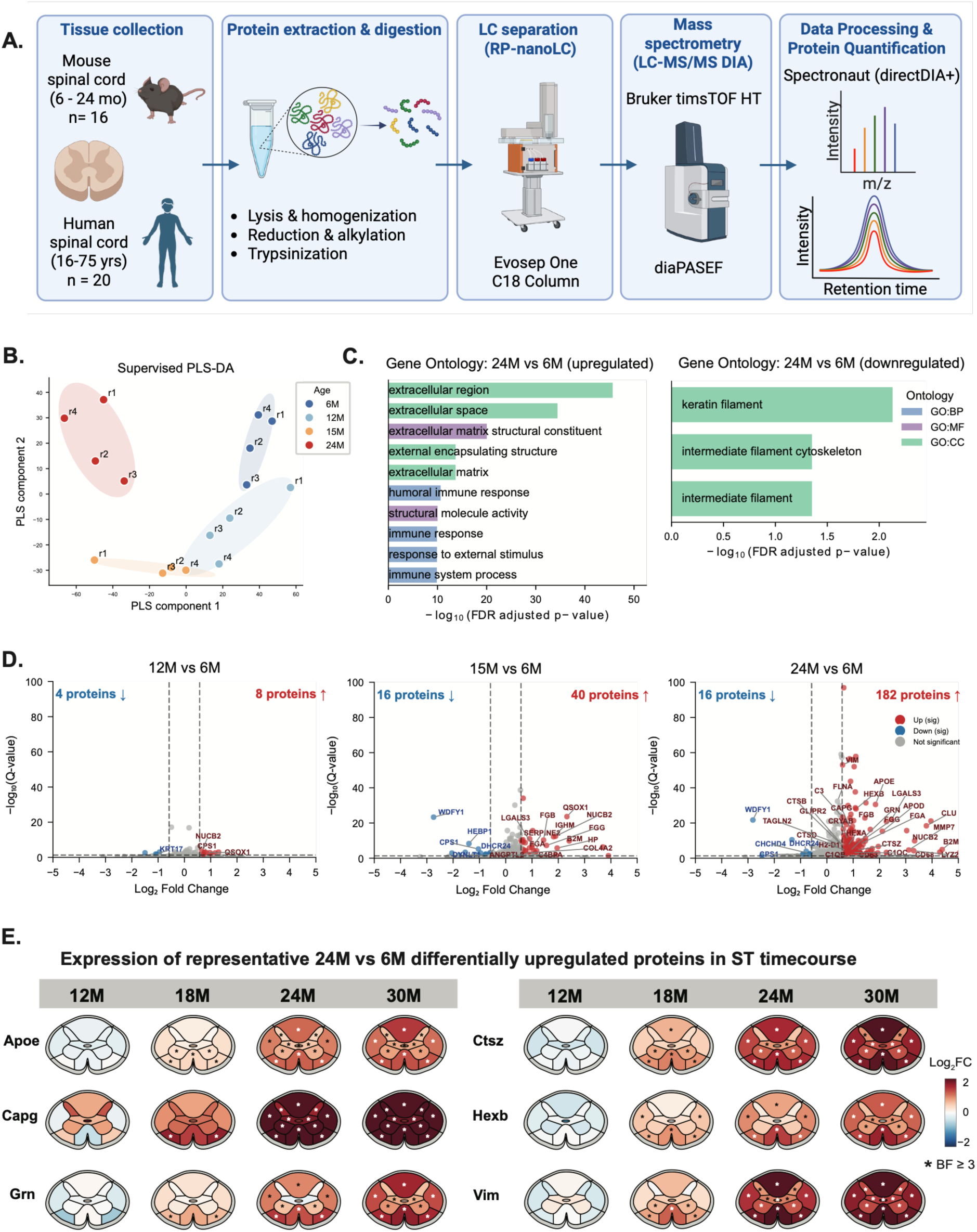
Global age-associated proteomic changes in the mouse spinal cord. **A)** Overview of workflow for LC-MS/MS for proteomic evaluation of mouse and human spinal cord. Mouse and human non-neuropathological spinal cord were collected, and protein was extracted and digested to tryptic peptides. Parallel accumulation-serial fragmentation combined with data-independent acquisition (diaPASEF) was used for data acquisition, and protein quantified for downstream analysis. **B)** Multigroup partial least squares-discriminant analysis (PLS-DA) comparing spinal cord from 6-, 12-, 15-, and 24-month mice (n=4 mice/group). Each dot represents one mouse, and ellipses show group dispersion in latent variable space. **C)** Gene Ontology (GO) terms for significantly altered proteins upregulated and downregulated at 24 months vs. 6 months in mouse spinal cord. **D)** Volcano plots for differentially abundant proteins at 12 months, 15 months, and 24 months vs. 6 months. (log2FC > 0.58 (∼1.5 fold), q-value < 0.05). Increasing proteins are shown in red, decreasing proteins are shown in blue, and proteins not meeting thresholds for significance are shown in grey. **E)** Spatial gene expression changes of six representative differentially upregulated proteins at 24 months across mouse ST time course. Color indicates log2 fold change of each gene at 12-, 18-, 24-, and 30-month time points relative to 6 months across spinal cord regions, and asterisk indicates significant difference between time point and 6 months (BF ≥ 3).

To examine the age-associated variation in the global protein abundance profiles, we performed partial least squares discriminant analysis (PLS-DA). This supervised analysis separated samples by age group, consistent with age-related differences in their overall proteomic profiles (Figure 2B). We next performed differential protein abundance analysis comparing mice at 12, 15, and 24 months of age to 6-month-old mice. Differentially abundant proteins (DEPs) were defined by an absolute log_2_ fold change > 0.58 and an FDR-adjusted p-value (q-value) < 0.05. Gene Ontology (GO) enrichment analysis of DEPs identified at 24 months revealed that upregulated proteins were predominantly associated with the extracellular matrix (ECM) and immune-related processes, whereas downregulated proteins were enriched for processes related to keratin filaments and the intermediate filament cytoskeleton (Figure 2C). Examination of differential protein abundance across the aging time course revealed an increasing number of DEPs, with the greatest number observed at 24 months (Figure 2D). Notably, the majority of these changes reflected an increased abundance of proteins associated with inflammation and immune response (e.g. CD68, C1QC, APOE), lysosomal function (e.g. LGALS3, HEXB), and ECM remodeling (e.g. MMP7, VIM), whereas relatively few proteins exhibited decreased abundance.

We next examined whether, when, and where these proteomic alterations were apparent at the transcriptional level. We compared proteins upregulated at 24 months with differentially expressed genes (DEGs) identified in our mouse spatial transcriptomics (ST) dataset (Bayes Factor ≥ 3 defines a DEG in each anatomical region; described in detail below and in Methods). Approximately 40% (72/182) of the upregulated proteins exhibited corresponding transcriptional upregulation in at least one anatomical region and timepoint in the mouse ST dataset, indicating partial concordance between the aging changes in protein and transcript abundance. Interestingly, ST analysis revealed that a subset of genes corresponding to proteins first upregulated at 24 months already exhibited localized transcriptional upregulation at 18 months, predominantly within the ventral white matter. By 24 and 30 months, these transcriptional changes were observed more broadly across white matter regions and extended into the grey matter, suggesting a spatial expansion of age-associated molecular alterations (Figure 2E). This spatiotemporal pattern was observed for genes including *Apoe*, *Capg*, *Grn*, *Ctsz*, *Hexb*, and *Vim*, which are associated with lipid metabolism, lysosomal function, immune responses, and cytoskeletal organization. We used immunofluorescence protein imaging to validate this, and we found a similar spatiotemporal pattern in APOE expression (Figure S4, Table S13). Together, these findings suggest that a subset of age-associated proteomic changes is accompanied by earlier, regionally localized transcriptional alterations, highlighting the spatial heterogeneity of molecular changes during spinal cord aging.

Next, we sought to determine whether similar age-associated proteomic alterations occur in the human spinal cord. We analyzed post-mortem lumbar spinal cord tissue from 20 non-neurological control donors aged 16-75 years (Figure 2A). Donors were stratified into young (16-34 years), middle-aged (47-60 years) and older (62-75 years) groups to examine changes in protein abundance across adulthood. Using LC-MS/MS, we quantified 6,564 proteins across the human spinal cord samples and assessed differential protein abundance between age groups (Table S6). The PLS-DA analysis showed separation of samples across the three age groups (Figure 3A). Differential protein abundance analysis was performed between young, middle-aged, and older donors, using an absolute log_2_ fold change > 0.58 and an FDR-adjusted p-value (q-value) < 0.01 to define DEPs. GO enrichment analysis comparing old and young donors revealed that upregulated proteins were predominantly associated with the fibrinogen complex, cell adhesion, vasoconstriction, blood coagulation, and basement membrane organization, whereas downregulated proteins were enriched for neurofilament organization, intermediate filament assembly, and collagen-associated structures (Figure 3B).

**Figure 3.**
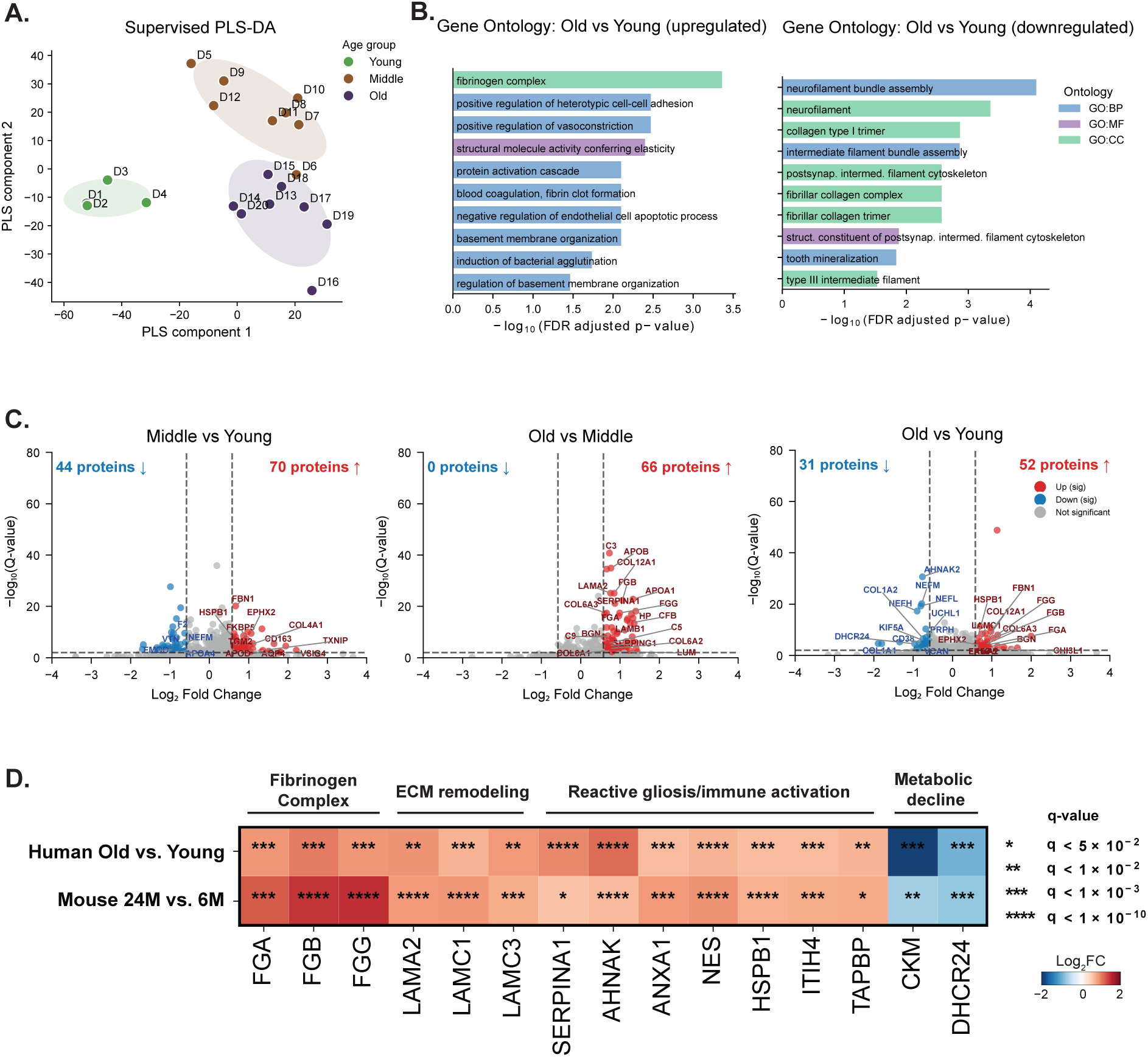
Global age-associated proteomic changes in the human spinal cord. **A)** Multigroup partial least squares-discriminant analysis (PLS-DA) comparing spinal cord from young (16-34 years, n=4), middle (47-60 years, n=8), and old age (62-75 years, n=8) human donors. Each dot represents one donor, and ellipses show group dispersion in latent variable space. **B)** Gene Ontology (GO) terms for significantly altered proteins upregulated and downregulated in Old vs. Young human spinal cord. **C)** Volcano plots for differentially abundant proteins in Middle vs. Young, Old vs. Middle, and Old vs. Young groups (log2FC > 0.58 (∼1.5 fold), q-value < 0.01). Increasing proteins are shown in red, decreasing proteins are shown in blue, and proteins not meeting thresholds for significance are shown in grey. **D)** Heatmap of shared significantly altered proteins between Old vs. Young human spinal cord and 24 month vs. 6 month mouse spinal cord comparisons. Color indicates log2FC, and asterisk indicates q-value.

Examination of differential protein abundance across age groups revealed distinct patterns of proteomic alterations (Figure 3C). Relative to young donors, middle-aged donors exhibited 70 upregulated and 44 downregulated proteins. In contrast, the comparison between older and middle-aged donors identified 66 upregulated proteins and no significantly downregulated proteins, suggesting that detectable proteomic changes during this later aging interval were predominantly characterized by increased protein abundance. Comparison of old and young donors identified 52 upregulated and 31 downregulated proteins. Notably, several proteins associated with extracellular matrix organization and vascular-related processes, including FBN1, FGG, FGB, and COL6A3 were increased in older donors, while proteins associated with neuronal cytoskeletal organization, including NEFM and NEFH, were reduced in abundance. Together, these findings highlight age-associated alterations in extracellular matrix, vascular-associated and neuronal structural proteins in the human spinal cord.

Finally, we assessed whether there were shared proteomic alterations between mouse and human spinal cord aging, by comparing the DEPs in 24-month-old mice relative to 6-month-old mice with those identified in older human donors relative to young donors. We identified several proteins exhibiting concordant changes in abundance across species (Figure 3D). Notably, all three fibrinogen chains (FGA, FGB, and FGG) were significantly upregulated in both mice and humans, alongside proteins associated with ECM remodeling (LAMA2, LAMC1, and LAMC3) and reactive gliosis and immune-associated processes (SERPINA1, AHNAK, ANXA1, NES, HSPB1, ITIH4, and TAPBP). Conversely, CKM and DHCR24, which are implicated in energy metabolism and cholesterol biosynthesis, respectively, exhibited decreased abundance in both species with age. Overall, these findings identify shared age-associated proteomic alterations in mouse and human spinal cords.

### Spatiotemporally coordinated gene programs in the aging mouse spinal cord

Next, we wanted to examine broader spatiotemporal changes in aging at the gene expression level, given the distinct cellular composition encompassing grey matter and white matter architecture in the spinal cord. To study how aging affects gene expression patterns in the spinal cord, we fit Splotch, a Bayesian hierarchical model to our Visium spatial transcriptomics data (Figure 4A)^46,47,54–57^. Splotch models gene expression at each spatial spot as a zero-inflated Poisson distribution, with the spot-level expression rate λ determined by spot- and donor-level metadata (spinal cord region and age), spatial autocorrelation among neighboring spots, and spot-specific noise (Methods).

**Figure 4.**
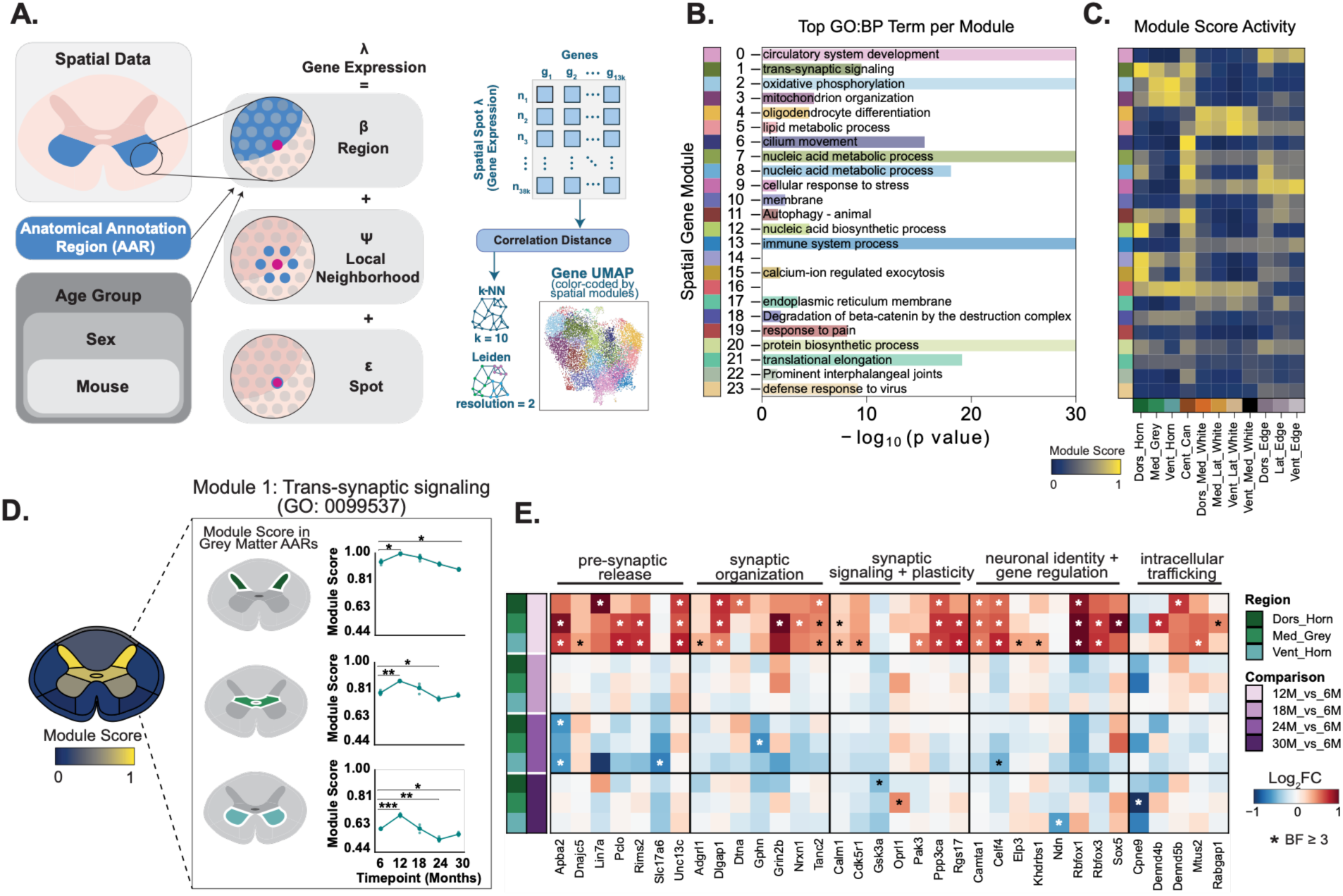
Spatiotemporally coordinated gene programs in the aging mouse spinal cord. **A)** The Splotch model for ST data fits spot-level gene expression (λ) on each array as a combination of mouse age group, sex, individual mouse, and specific region of interest (β), a spatial autoregressive component (ψ), and a spot noise (ε) component. Correlation distance between genes is calculated from modeled spot-level gene expression (λ) across all gene pairs across all spots. Genes are embedded into a k-Nearest Neighbors (k-NN) graph, and Leiden community clustering algorithm is used to identify spatially correlated gene modules. Each dot in the UMAP represents a gene colored by spatial module. **B)** Most significant Biological Process (BP) GO terms for each spatial gene module. Bar plot length indicates -log10(p-value). Empty rows indicate no enriched term. **C)** Average module scores for each spinal cord region. **D)** M1 (Trans-synaptic signaling) module score on an anatomical heatmap of the spinal cord. Line plots depict changes in module score in dorsal horn, medial grey, and ventral horn across time course. Asterisks indicate significant differences at time point compared to 6 months (Welch’s t-test, Benjamini-Yekutieli correction p < 0.05). **E)** Heatmap of M1 genes in mouse ST data across time course in dorsal horn, medial grey, and ventral horn. Each row represents a spinal cord region, and each column represents a gene. Color indicates log2 fold change of each gene at 12-, 18-, 24-, and 30-month time points relative to 6 months, and asterisk indicates significant difference between time point and 6 months (BF ≥ 3).

To identify gene programs that exhibit spatially coordinated expression, we grouped genes into spatial co-expression modules based on pairwise Pearson correlations calculated from Splotch-modeled spatial transcriptomics data. We then embedded each gene transcript into a k-nearest neighbor (k-NN) graph using Pearson correlation distance and performed Leiden clustering. This resulted in 24 spatiotemporal gene expression modules, each made up of genes with similar spatial expression patterns across the spinal cord (Figure 4A, Table S7,8). To investigate the biological functions underlying these spatial co-expression modules, we performed Gene Ontology (GO) enrichment analysis to identify biological processes overrepresented among the genes within each module (Figure 4B, Table S9). For instance, module 0 (M0) is linked to circulatory system development, M13 is linked to immune system processes, and M15 is linked to calcium-ion regulated exocytosis.

To characterize the spatial distribution and cell-type associations of these transcriptional programs, we calculated a module score for each of the 24 modules, defined as the weighted average expression of a module’s constituent genes (Table S8). Module scores were calculated across each individual spatial spot and across each nucleus in the orthogonal mouse spinal cord snRNA-seq dataset. These analyses revealed distinct anatomical and cellular expression patterns across modules (Figure 4C, Figure S5A,B). For example, M4 (oligodendrocyte differentiation), showed elevated expression in the white matter and highest expression in oligodendrocytes, while M1 (trans-synaptic signaling) was most highly expressed in the spinal cord grey matter and within neuronal cell populations including excitatory, inhibitory and motor neurons (Figure S5A,B). These modules thus define region-and cell type-specific gene programs in the mouse spinal cord.

### Aging-associated molecular programs exhibit divergent trajectories across the mouse spinal cord grey and white matter

We wanted to understand whether these spatial modules exhibit distinct trajectories across the mouse lifespan. Given the differences in cellular composition and functional organization of the spinal cord grey and white matter, we also sought to characterize whether aging-associated spatiotemporal transcriptional changes differed between anatomical regions. We identified several modules exhibiting divergent age-associated trajectories, and these modules were enriched in different regions of the spinal cord and in different cell types (Figure 4D, 5, S5, Table S8).

We found a decrease in scores in multiple modules with age, including M1, M3, M11 and M15, and these modules tended to be more strongly enriched in the grey matter regions and primarily in neuronal populations. M1 (trans-synaptic signaling) showed an interesting spatiotemporal pattern, characterized by an increase at 12 months relative to 6 months before declining at later ages (Figure 4D). This module was highly enriched in the grey matter regions of the spinal cord and in neuronal populations. To investigate the genes driving the age-associated changes in the module scores, we performed differential gene expression analysis within each anatomical region, comparing mice aged 12, 18, 24, and 30 months to 6-month-old mice. For each gene, we quantified the evidence of an age-associated expression change using a Bayes Factor (BF), calculated using the Savage Dickey density ratio between the Splotch-derived posterior estimates of age contribution per anatomical region for that gene (Table S10,11). Genes with a BF ≥ 3 were classified as differentially expressed. At the gene level, the same temporal pattern was broadly reflected in the expression of M1 constituent genes involved in presynaptic release, synaptic organization and signaling, neuronal identity and intracellular trafficking (Figure 4E).

Module M3 (mitochondrion organization) was highly enriched in the grey matter and was most highly expressed in neuronal populations (Figure 5A). Its expression remained high over the time course and significantly declined at 30 months in the grey matter. Module M11 (autophagy) was also highly enriched in grey matter regions of the spinal cord and in neuronal populations, and its expression declined significantly at 30 months (Figure 5B). Finally, module M15 (calcium ion regulated exocytosis), was also most highly enriched in grey matter regions and in neuronal cells, particularly in the dorsal horn, and showed a significant decline with age in grey matter at 30 months (Figure 5C). While all of these modules also declined significantly at 30 months in the white matter, their overall expression was lower compared to the grey matter. The similar age-associated declines in M15 and M1 (trans-synaptic signaling), both enriched in neuronal populations and in grey matter regions, suggest coordinated downregulation of transcriptional programs supporting synaptic communication during spinal cord aging.

**Figure 5.**
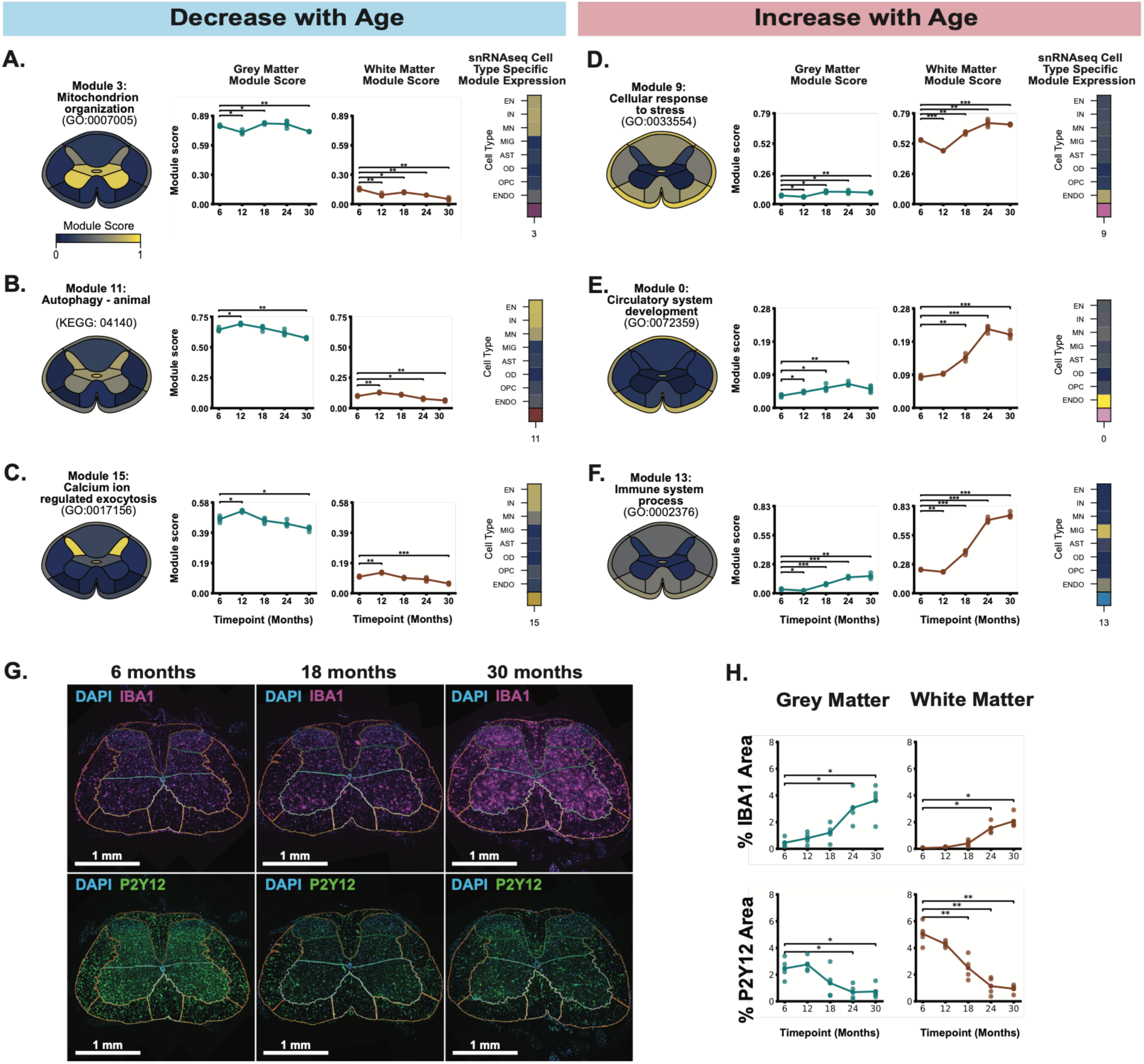
Aging-associated molecular programs exhibit diverging patterns of regulation in the mouse spinal cord grey and white matter. **A-C)** Average spatial gene module score per spinal cord region for modules declining with aging (left). Line plot depicting module score trends over mouse time course for grey matter (middle) and white matter (right) regions. Asterisks indicate significant differences at time point compared to 6 months (Welch’s t-test, Benjamini-Yekutieli correction p < 0.05). **D-F)** Average spatial gene module score per spinal cord region for modules increasing with aging (left). Line plot depicting module score trends over mouse time course for grey matter (middle) and white matter (right) regions. Asterisks indicate significant differences at time point compared to 6 months (Welch’s t-test, Benjamini-Yekutieli correction p < 0.05). **G)** Representative IF image for DAPI, IBA1, and P2Y12 staining at 6, 18, and 30 months. **H)** Mean IBA1 area as a percentage of total area in grey matter and white matter (top), and mean P2Y12 area as a percentage of total area in grey matter and white matter (bottom). Asterisks indicate significant differences at time point compared to 6 months (Welch’s t-test, Benjamini-Hochberg correction p < 0.05).

In contrast, several modules (e.g. M9, M0, M13) exhibited increased expression with advancing age, and these tended to be more highly enriched in white matter regions and associated with vascular and glial cell types. While the module scores generally increased in both grey matter and white matter with age, they increased to a much greater degree in the white matter. M9 (cellular response to stress) was highly enriched in the white matter and in endothelial cells. It was significantly upregulated in the white matter at 18, 24 and 30 months (Figure 5D). M0 (circulatory system development) was also enriched in the white matter and in the edges of the spinal cord, where the vascular cells reside (Figure 5E). Consistent with this, it was most highly expressed in endothelial cells, and this module was significantly upregulated at 18 months and at subsequent timepoints. M13 (immune system process) was also enriched in the white matter, and its expression was driven by microglia and increased significantly with each time point (Figure 5F).

Given the pronounced age-associated changes in the microglia-enriched M13 module, we sought to determine whether these transcriptional changes were accompanied by alterations in microglial populations at the protein level. To this end, we performed immunofluorescence (IF) staining for microglial markers IBA1 and P2Y12 in spinal cord tissue from mice aged 6 to 30 months (Figure 5G). IF showed an increase in IBA1+ area and cell counts with age in both grey and white matter regions, whereas P2Y12+ area decreased in both regions and P2Y12+ cell counts decreased in the white matter (Figure 5H, Figure S6; n=4 mice per age group). These changes reflect a shift in microglial populations toward a less homeostatic state, in line with the changes observed at the transcriptional level in M13. This shift in microglial states in the white matter also resembles changes to white matter microglia in the aging mouse brain^58^. Altogether, these results suggest region- and cell type-specific changes with age; modules that are enriched in neuronal populations in grey matter are more susceptible to decline in gene expression, and vascular and glial populations in white matter are more susceptible to increase in gene expression.

### Age-associated vascular and immune remodeling reveals enrichment of senescence-associated programs in mouse and human spinal cord

Given the role of cellular senescence as a key hallmark of aging contributing to age-related tissue dysfunction, we wanted to investigate whether any of the age-related changes in our ST modules were related to cellular senescence. We investigated whether any modules were enriched for senescence-associated signatures, including SASP, DNA damage response, and cell-cycle arrest. We selected four established senescence gene sets: Fridman Senescence Up^59^, San Diego Tissue Mapping Center (SD TMC), SenMayo^60^, and SenBrain^61^ (Figure 6A, Table S12). Although the genes in these lists are largely non-overlapping, M0 (circulatory system development) and M13 (immune system process) contained the highest proportions of genes from multiple lists (Figure 6B).

**Figure 6.**
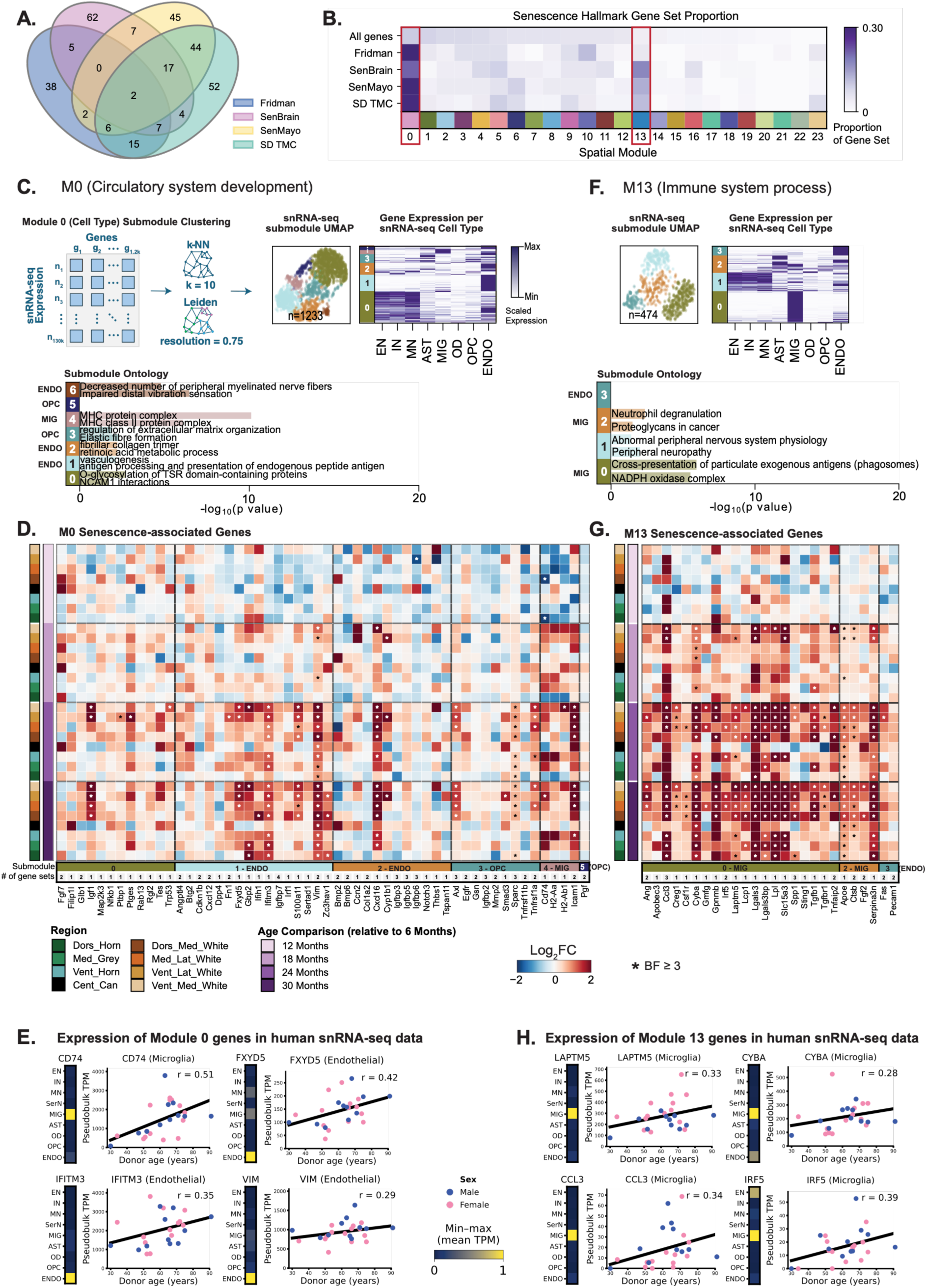
Age-associated vascular and immune remodeling reveals enrichment of senescence-associated programs in mouse and human spinal cord. **A)** Overview Venn diagram of externally curated senescence gene set lists used in this study. Numbers in intersecting sections indicate number of overlapping genes. SD TMC = San Diego Tissue Mapping Center. **B)** Senescence hallmark gene set proportions within each spatial module. “All genes” includes all genes assigned to a spatial module (n=13,280 genes). Red outline highlights modules with the highest proportion of senescence genes across all gene sets (M0 and M13). **C)** Representative schematic of submodule clustering. Genes from spatial module 0 are assigned into submodules using correlation with snRNA-seq data. M0 gene-gene k-NN is embedded into a UMAP, where each dot represents a gene that is colored according to the assigned submodule. Average gene expression in M0 for each snRNA-seq cell type, with expression values min-max scaled per gene for visualization, and submodule assignment annotated on the left. Submodule GO terms for M0 (bottom). Length of underlying bar plot indicates -log10(p-value). **D)** Heatmap of senescence-associated genes in M0 (Circulatory system development) across spinal cord regions and time points. Genes are split into their respective submodules and corresponding snRNA-seq cell type, and number of gene sets each gene is found in is listed. Color indicates log2 fold change of each gene at 12-, 18-, 24-, and 30-month time points relative to 6 months, and asterisk indicates significant difference between time point and 6 months (BF ≥ 3). **E)** Heatmap showing mean transcripts per million (TPM; min-max scaled for visualization) across cell types in human snRNA-seq data per gene in M0. Pseudobulk TPM plotted against age across donors in microglia (*CD74*, top left) and endothelial cells (*FXYD5*, top right; *IFITM3*, bottom left; *VIM*, bottom right). **F)** M13 (Immune system process) submodule UMAP, average gene expression for each snRNA-seq cell type, and submodule GO terms. **G)** Heatmap of senescence-associated genes in M13 across spinal cord regions and time points split into submodules and corresponding snRNA-seq cell type. Color indicates log2 fold change of each gene at 12-, 18-, 24-, and 30-month time points relative to 6 months, and asterisk indicates significant difference between time point and 6 months (BF ≥ 3). **H)** Heatmap showing mean transcripts per million (TPM; min-max scaled for visualization) across cell types in human snRNA-seq data per gene in M13. Pseudobulk TPM plotted against age across donors in microglia (*LAPTM5*, top left; *CYBA*, top right; *CCL3*, bottom left; *IRF5*, bottom right).

As previously established, M0 was most highly expressed in endothelial cells and exhibited a greater increase in expression with age in the white matter. To further resolve the transcriptional programs underlying M0, we subclustered its constituent genes into submodules using correlation distance between each gene pair in our orthogonal snRNA-seq spinal cord dataset, resulting in seven submodules with distinct expression patterns across cell types (Figure 6C, Figure S7). For instance, M0-1, M0-2, and M0-6 were submodules expressed primarily in endothelial cells, while M0-3 and M0-5 were expressed primarily in oligodendrocyte progenitor cells (OPCs), and M0-4 was expressed in microglia. GO enrichment analysis revealed that the genes in M0-1 were enriched for biological processes associated with vasculogenesis, whereas M0-2 and M0-3 were associated with fibrillar collagen trimer and regulation of extracellular matrix (ECM) organization, respectively. These findings suggest that M0 encompasses distinct transcriptional programs involved in vascular development and ECM remodeling, processes that may contribute to senescence-associated tissue alterations during spinal cord aging.

The 54 senescence-associated genes within M0 were primarily expressed in the endothelial M0-1 and M0-2, OPC-driven M0-3 and M0-5, and microglial M0-4 (Figure 6D). Notably, several senescence associated genes within these submodules were significantly upregulated with age, including interferon-response genes (*Gbp2, Ifih1, Ifitm3, Zc3hav1*), inflammatory and cell adhesion-associated genes (*Cxcl16, Icam1, Axl*) and genes involved in ECM remodeling (*Sparc, Vim*). These findings suggest that the senescence-associated signature within M0 is characterized predominantly by inflammatory and tissue-remodeling processes, consistent with inflammatory, SASP-like features.

To assess whether these transcriptional patterns are conserved during human spinal cord aging, we measured the expression of these genes in our aging human lumbar spinal cord snRNA-seq dataset. Notably, *CD74*, an antigen-presenting gene belonging to the microglia-associated M0-4 submodule in the mouse ST data, was also most highly expressed in human microglia. *CD74* expression exhibited a positive, albeit nonsignificant, correlation with donor age (Figure 6E, r = 0.51). The cell-adhesion gene *FXYD5*, the interferon-signaling gene *IFITM3*, and the cytoskeletal organization gene *VIM* (all belonging to the endothelial-associated M0-1), were each most highly expressed in human endothelial cells. The expression of each of these genes trended up with donor age (*FXYD5*, r = 0.42; *IFITM3*, r = 0.35; and *VIM*, r = 0.29; Figure 6E).

In addition to M0, the immune-associated M13 was also enriched for genes from the curated senescence gene sets. M13 was further subdivided into four submodules, including two microglia-enriched submodules associated with immune-related processes such as neutrophil degranulation and antigen presentation, and one endothelial-enriched module (Figure 6F). The 26 senescence-associated genes were predominantly expressed in the microglial M13-0 and M13-2, with significant age-associated upregulation emerging in the white matter at 18 months and subsequently in the grey matter at 24 and 30 months (Figure 6G). Further examination of M13 revealed increased expression of genes implicated in microglial activation (*Gpnmb, Spp1, Lgals3*), lipid metabolism (*Apoe, Lpl*), lysosomal function (*Ctsb, Laptm5*) and inflammatory signaling (*Ccl3, Irf5, Cyba*). Notably, several of these genes have previously been linked to disease-associated and white matter-associated microglial states^61,62^, suggesting that age-related increases in M13 expression may reflect a shift toward reactive microglial programs involved in lipid metabolism, clearance of cellular debris, and inflammatory responses.

Many of these genes were also most strongly expressed in the human microglia in our human snRNA-seq data. Among them were *LAPTM5*, which is involved in lysosomal trafficking; *CYBA*, encoding a component of the NADPH oxidase complex; *CCL3*, a pro-inflammatory chemokine; and *IRF5*, a transcriptional regulator of the innate immune response. Although none of these genes exhibited statistically significant age-associated changes, all four genes showed positive correlation with donor age in human microglia (*LAPTM5*, r = 0.33; *CYBA*, r = 0.28; *CCL3*, r = 0.34, *IRF5*, r = 0.39; Figure 6H). These trends are consistent with the mouse findings and suggest that inflammatory, oxidative, and endolysosomal transcriptional programs may also increase during human spinal cord aging.

### Enrichment of p21/*CDKN1A* and p16/*CDKN2A* in differing cell types in the human spinal cord

Because our transcriptional data revealed enrichment of senescence-associated signatures in vascular and microglial gene programs, we next asked whether these cell types were also enriched for the canonical senescence markers *CDKN1A*/p21 and *CDKN2A*/p16 in the human spinal cord. In the human snRNA-seq data, we found that *CDKN1A* was enriched most abundantly in endothelial cells and microglia (Figure 7A). In endothelial cells, *CDKN1A* expression was higher in female donors compared to male donors regardless of age (Figure 7Aiii, Welch’s t-test, p = 0.00256). Consistent with the transcriptional data, we stained for p21 and found positive staining in vascular cells (endothelial and smooth muscle cells) and microglia (Figure 7B).

**Figure 7.**
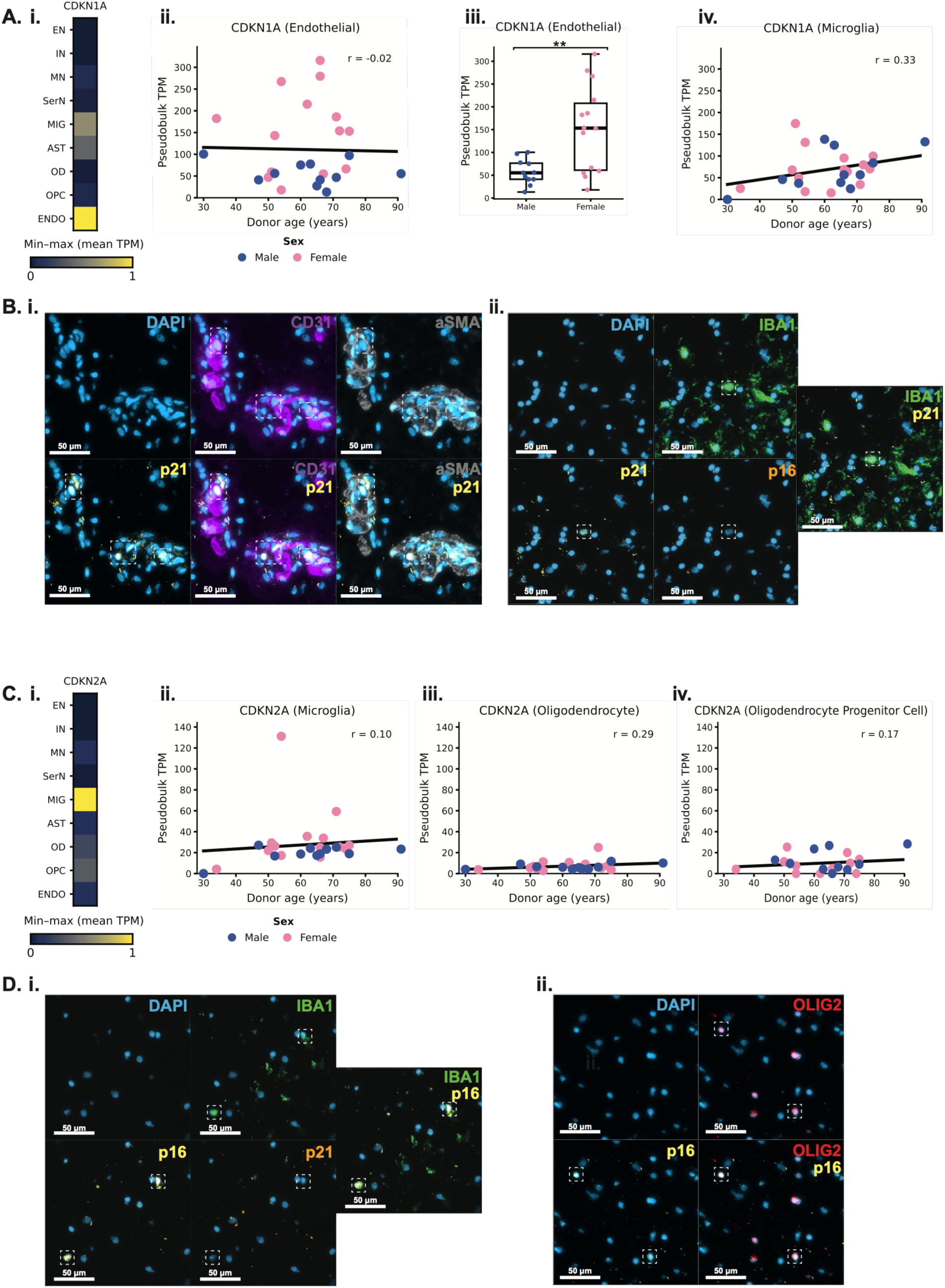
Enrichment of p21/*CDKN1A* and p16/*CDKN2A* in differing cell types in the human spinal cord. **A)** i) Heatmap showing *CDKN1A* mean transcripts per million (TPM; min-max scaled for visualization) across cell types in human snRNA-seq data. ii) Pseudobulk TPM plotted against age across donors in endothelial cells. iii) Pseudobulk TPM in male vs. female donors in endothelial cells (Welch’s t-test, p = 0.00256). iv) Pseudobulk TPM plotted against age across donors in microglia. **B)** Representative IF images of p21 with i) vascular markers (CD31 and aSMA) and ii) microglia (IBA1) in human spinal cord from donor NY7274. **C)** i) Heatmap showing *CDKN2A* mean TPM (min-max scaled for visualization) across cell types in snRNA-seq data. Pseudobulk TPM plotted against age across donors in ii) microglia, iii) oligodendrocytes, and iv) oligodendrocyte progenitor cells. **D)** Representative IF images of p16 with i) microglia (IBA1) and ii) oligodendrocytes (OLIG2) in human spinal cord from donor NY7341.

In contrast, *CDKN2A* expression was most abundantly enriched in microglia, followed by oligodendrocyte lineage cells (oligodendrocyte progenitor cells and oligodendrocytes) (Figure 7C). There were no sex differences in the abundance of *CDKN2A* transcript or age-related differences detected in our analysis. We also stained for p16 and found positive staining in microglia and oligodendrocyte cells (Figure 7D). While *CDKN1A* and *CDKN2A* were both enriched in microglia, IBA1+ microglia did not appear to express both markers at once (Figure 7Bii, 7Di). Overall, these results suggest potentially distinct, cell type-specific phenotypes between p16+ and p21+ cells.

## Discussion

In this study, we uncover converging aging and senescence-associated signatures across the mouse and human spinal cord, integrating LC-MS/MS, ST, snRNA-seq, and multiplexed protein imaging techniques. To our knowledge, our study also includes the largest non-neuropathological, aging human spinal cord snRNA-seq dataset to date.

At the global proteomic level, we found an increase with age in abundance of proteins associated with fibrinogen complex, ECM remodeling, and immune response in both mouse and human spinal cord. Intriguingly, fibrinogen is not usually expressed in the central nervous system (CNS), and its expression in the CNS is thought to be related to blood-brain barrier disruption^63,64^. The co-occurrence of fibrinogen and basement membrane laminin accumulation in both species, together with the increase in vascular module M0 in mouse white matter, suggests that age-associated changes in the vasculature and potentially the blood-spinal cord barrier may contribute to this early vulnerability. This is in line with prior evidence of blood-spinal cord permeability increasing in aged mice^4^.

Using the spatial co-expression patterns of genes from our mouse ST data, we derived ‘spatial modules’ to understand which biological processes change with age and where these changes are localized. Our data revealed several processes that decrease in expression with age, including synaptic signaling, mitochondrial organization, autophagy, and calcium ion regulated exocytosis. These gene programs were primarily enriched in the grey matter, and module scoring in reference snRNA-seq datasets revealed that they were primarily found in excitatory, inhibitory, and motor neurons. Prior studies have also shown changes in aging motor neurons related to synaptic pruning and oxidative stress^2,65^. While we saw a significant decline in these modules in grey matter at 30 months, the overall gene expression remained relatively high even in advanced age. This could be associated with high resilience in neuronal populations in the spinal cord, in line with the critical roles that these cells play in processes such as movement, respiration, autonomic regulation, and sensation.

In contrast, increasing spatial modules in the spinal cord were mainly vascular and immune in nature. Genes associated with these modules were primarily enriched in the white matter, and their module scores increased to a greater degree in white matter compared to grey matter. Furthermore, these modules were enriched for senescence-associated genes from the SenMayo, SenBrain, Fridman, and SD TMC gene set lists. Many of the genes in these modules were enriched in endothelial and microglial populations in both the mouse and human spinal cord. They increased significantly in the mouse microglial- and endothelial-specific spatial submodules and trended up in the same cell populations in the human snRNA-seq data with age, suggesting convergent aging and senescence-related signatures in the vascular and microglial cell types across species. In the mouse, the progressive loss of the homeostatic marker P2Y12 alongside increased IBA1, concentrated in the white matter, further supports a shift of white matter microglia toward more reactive states^44,58,66^. Our findings that the white matter in the spinal cord is a hotspot for senescence-associated aging changes overlap with several recent studies that have similarly identified the white matter as a hotspot for senescence in the mouse and human brain (cortex and hippocampus)^44,47,58,67^. This suggests that the white matter may be particularly vulnerable to aging and cellular senescence in the central nervous system, regardless of region or species.

Finally, using *CDKN1A*/p21 and *CDKN2A*/p16 as proxies for senescent cells, in the human spinal cord, we found enrichment for *CDKN1A*/p21 in endothelial cells and microglia, and *CDKN2A*/p16 in microglia and oligodendrocyte lineage cells. These findings overlap with cell-type specific p21 and p16 expression in the human cortex and hippocampus^47,67^. Furthermore, *CDKN1A* transcripts in endothelial cells were more abundantly expressed in the spinal cords of females, which was a trend also seen in the human hippocampus^47^. While neither of these genes were significantly associated with age, both genes exhibited a positive, albeit non-significant, correlation with age in specific cell types (*CDKN1A* in microglia, *CDKN2A* in oligodendrocytes). Quantification of our protein imaging will 1) provide orthogonal validation for these age-and sex-associated trends, 2) reveal whether p16+ and p21+ microglia represent overlapping or distinct populations, and 3) show how the distribution of these cells vary across grey vs. white matter regions.

Here, we identify microglia and endothelial cells as populations vulnerable to aging across both mouse and human spinal cord. Future studies should examine whether and how these cell types and their senescence-associated signatures may be implicated in spinal cord diseases, including spinal cord injury and amyotrophic lateral sclerosis (ALS). Altogether, this work represents the first comprehensive, multimodal mapping of aging and senescence across the mouse and human spinal cord.

## Limitations

While our study now includes the largest non-neuropathological, aging human spinal cord snRNA-seq dataset, we were still limited in power by the small number of young donors. This is attributed to challenges with sourcing postmortem human spinal cord tissue, as this region is not often banked, and the majority of donors included were prospectively banked specifically for this study. Our mouse ST and IF data was also limited to two male and two female mice per timepoint, so we were not powered to detect sex differences.

Technical limitations include the limited resolution of bulk mass spectrometry data. Given the lack of spatial resolution, we aimed to circumvent this limitation by mapping the spatial location of differentially abundant proteins onto our mouse ST data. While our ST technology did not provide single-cell resolution to define exact cell subpopulations and interactions driving spatial gene programs, next generation spatial technologies will hopefully provide higher resolution to resolve these interactions in future studies.

## Materials and Methods

### TISSUE ACQUISITION

#### Mice

The C57Bl/6J (strain #000664) mouse lines used for spatial transcriptomics and immunofluorescence experiments were obtained from the Jackson Laboratory and aged in the lab of Dr. Neil Shneider. Aged C57BL/6NCrl (strain #701) male mice (6, 12, 15 and 24-months) used for mass spectrometry profiling were obtained from Charles River Laboratories. All animals were maintained in a full-barrier, pathogen-free facility with an ambient temperature of 18-23 °C, 40-60% humidity, and a 12-hour light/dark cycle, in accordance with ethical guidelines established and maintained by Columbia University Irving Medical Center’s Institutional Animal Care and Use Committee.

#### Mouse Spinal Cord Collections

Mice were transcardially perfused with pre-chilled 1X Phosphate Buffered Saline (PBS), and spinal cords were subsequently dissected. The L3-L5 lumbar region was isolated based on ventral root anatomy and embedded in Tissue Plus OCT Compound. Tissue molds were flash frozen in a bath of dry ice and pre-chilled ethanol and stored at -80°C.

#### Post-mortem Human Tissue

Human post-mortem lumbar spinal cord tissue from sudden death non-neurological control donors was acquired through New York Brain Bank (NYBB) at Columbia University, Georgetown, University of California San Diego, and Yadav et al^68^. Samples were collected in accordance with institutional review board (IRB) protocols, and were transferred to Columbia University Senescence Tissue Mapping Center (CUSTMAP) following all applicable regulations for processing, sequencing, and analysis. Metadata for the donors included in this study can be found in Table S1.

### LIQUID CHROMATOGRAPHY/MASS SPECTROMETRY (LC-MS/MS)

#### Cohorts for Proteomic Analysis and Tissue Lysis and Homogenization

Human lumbar spinal cord tissue was obtained from 20 participants ranging from 16 to 75 years of age with 4 young adults (16-34 years old), 8 middle-aged adults (47-60 years old) and 8 elderly adults (62-75 years old). The mouse experimental paradigm contained four biological replicates (n=4 per group) in each of the 6-, 12-, 15-, and 24-month age groups. All samples were prepared for proteomic analysis (data-independent acquisition, DIA)^69,70^ as follows: samples were homogenized in ∼300 µL of lysis buffer containing 8 M urea, 2% sodium dodecyl sulfate (SDS), 1 µM trichostatin A (TSA), 3 mM nicotinamide adenine dinucleotide (NAD), 75 mM sodium chloride, and 1x protease and phosphatase inhibitor cocktail (Thermo Fisher Scientific, Waltham, MA) in 200 mM triethylammonium bicarbonate (TEAB) in 2.0 mL safe-lock tubes (VWR International, Radnor, PA) containing stainless steel beads with three 30-second intervals of shaking at 25 Hz using a Qiagen TissueLyser II (Qiagen, Hilden, Germany). Tissue homogenates were centrifuged at 15,700 x g for 10 min at 4°C, and the supernatant was collected for label-free quantitative proteomics experiments. Protein concentration was determined using a bicinchoninic acid (BCA) assay (Thermo Fisher Scientific, Waltham, MA).

#### Protein Digestion and Desalting

Aliquots containing 100 µg of human protein or 200 µg of mouse protein were adjusted to 50 µL with water, reduced using 20 mM dithiothreitol in 50 mM TEAB at 50°C for 10 min, cooled to room temperature (RT), held at RT for 10 min, and alkylated using 40 mM iodoacetamide in 50 mM TEAB at RT in the dark for 30 min. Samples were acidified with 12% phosphoric acid to obtain a final concentration of 1.2% phosphoric acid. S-Trap buffer, consisting of 90% methanol in 100 mM TEAB at pH ∼7.1, was added, and samples were loaded onto the S-Trap micro spin columns^71^. The entire sample volume was spun through the S-Trap micro spin columns at 4,000 x g and RT, binding the proteins to the micro-spin columns. Subsequently, S-Trap micro spin columns were washed twice with S-Trap buffer at 4,000 x g and RT and placed into clean elution tubes. Samples were incubated for one hour at 47°C with sequencing-grade trypsin (Promega, San Luis Obispo, CA) dissolved in 50 mM TEAB at a 1:25 (w/w) enzyme-to-protein ratio. An additional aliquot of trypsin dissolved in 50 mM TEAB was added, and samples were digested overnight at 37°C. Peptides were sequentially eluted from S-Trap micro spin columns with 50 mM TEAB, 0.5% formic acid (FA) in water, and 50% acetonitrile (ACN) in 0.5% FA. After centrifugal evaporation, samples were resuspended in 0.2% FA in water and desalted with Oasis 10-mg Sorbent Cartridges (Waters, Milford, MA). The desalted elutions were subjected to centrifugal evaporation and resuspended in 0.1% FA in water at a final concentration of 1 µg/µL.

#### Mass Spectrometric Analysis

Both the mouse and human samples (400 ng of peptides per sample) were loaded onto EvoTips Pure (Evosep, Odense, Denmark) following the manufacturer’s protocol. During sample loading onto EvoTips Pure, one microliter of indexed retention time standard (iRT, Biognosys, Schlieren, Switzerland) was added to each sample. HPLC-MS/MS analyses were performed on an Evosep One liquid chromatography system (Evosep) coupled to a timsTOF HT mass spectrometer (Bruker, Bremen, Germany). The solvent system consisted of 0.1% FA in water (solvent A) and 0.1% FA in ACN (solvent B). Peptides were eluted on a PepSep C_18_ analytical column (150 µm x 15 cm, 1.5 µm particle size; Bruker) using the 30 samples per day (SPD) method^72^ with a 44-minute gradient length (500-nL/min flow rate). A zero-dead volume emitter was installed in the nano-electrospray source (CaptiveSpray source, Bruker Daltonics), and the source parameters were set as follows: capillary voltage 1600 V, dry gas 3 L/min, and dry temperature 180°C. Each sample was acquired in data-independent acquisition coupled to parallel accumulation-serial fragmentation (diaPASEF) mode^73^ with 1 survey TIMS MS scan and 24 diaPASEF MS/MS scans per 2.03-s cycle. The dual TIMS analyzer was operated at a 100% duty cycle, with equal accumulation and ramp times of 75 ms each. For each TIMS MS scan, the ion mobility ranged from 1/K_0_ = 0.85-1.30 Vs/cm^2^, and the mass range covered *m/z* 100-1,700. For both datasets, the diaPASEF acquisition scheme comprised 63 contiguous 15-Th isolation windows spanning *m/z* 305.86-1,250.9, covering an inverse ion mobility range of 0.74-1.30 Vs/cm^2^. The collision energy was defined as a linear function of mobility, starting from 20 eV at 1/K_0_ = 0.6 Vs/cm^2^ to 59 eV at 1/K_0_ = 1.6 Vs/cm^2^. For calibration of the ion mobility dimension, three ions of Agilent ESI-Low Tuning Mix were selected (m/z, 1/K0[Vs/cm^2^]: 622.0289, 0.9915; 922.0097, 1.1986; 1221.9906, 1.13934).

#### DIA Data Processing and Statistical Analysis

Mouse and human diaPASEF data were processed separately in Spectronaut v19.0.240606.62635 and v20.4.260109.92449, respectively, using directDIA+^74^. Mouse data were searched against uniprot_mouse_proteome.fasta (58,430 entries), and human data against uniprotkb_human_proteome_UP000005640_reviewed-only_2026_02_10.fasta (20,416 entries). Searches used specific Trypsin/P digestion, up to two missed cleavages, and peptide lengths of 7-52 amino acids. Cysteine carbamidomethylation was a fixed modification; methionine oxidation and protein N-terminal acetylation were variable modifications, with a maximum of five variable modifications. Pulsar search false discovery rate thresholds were 1% at the peptide-spectrum match, peptide, and protein-group levels. DIA precursor and experiment-level protein q-value cutoffs were 0.01; the run-level protein q-value cutoff was 0.05. Dynamic extraction windows and mass tolerances, precision iRT^75^ with local non-linear calibration, and iRT profiling were enabled. Quantification used MS2 peak areas, intensity-based selection of 3-6 fragments per peptide, interference correction, and local cross-run normalization with sparse row selection (identified in at least one run). Precursors were filtered by identification q-value, with background-signal imputation enabled. Protein inference used IDPicker; protein quantities were calculated by summing peptide quantities, with up to seven peptides per protein group and up to ten precursors per stripped peptide sequence. The protein LFQ method was QUANT 2.0 for mouse data and Automatic for human data. The mouse experimental paradigm contained four biological replicates (n=4 and correspondingly four acquisitions) in each of the 6-, 12-, 15-, and 24-month age groups, and unpaired t-tests were performed. The human experimental paradigm compared 12 individuals aged 60 years or younger with eight individuals older than 60 years old using unpaired t-tests without assuming equal variance. Both analyses were performed with group-wise testing corrections using the Storey method^76^ and the following false discovery rates were used for quantification candidate-filter settings: mouse study q<0.05, human study q<0.01 (a more stringent threshold was applied to the human cohort to account for greater inter-individual heterogeneity and uncontrolled confounding factors) and an absolute log2 ratio threshold of >0.58.

### VISIUM SPATIAL TRANSCRIPTOMICS (ST)

#### Tissue Processing, RNA Capture, and Library Preparation for Spatial Transcriptomics

Fresh frozen spinal cord tissue blocks were equilibrated to -16°C for 30 minutes in the cryostat before sectioning. Cryosections were cut at 10 µm thickness as serial adjacent sections from the same tissue block onto pre-chilled Visium Spatial Gene Expression slides. Each array comprised four technical replicates per mouse, and each Visium slide included sections from two male and two female mice within the same age group to minimize technical batch effects. Slides were stored at -80°C for a maximum of three days before processing.

Spatially resolved gene expression data was generated following the guidelines outlined in the 10X Genomics Visium Spatial Gene Expression Reagent User Guide (CG000239, Rev F). Briefly, tissue sections were fixed in methanol at -20°C for 30 minutes, and stained with hematoxylin and eosin (H&E). Brightfield RGB histological images were acquired using the Zeiss Axio Observer Z1 microscope equipped with an EC Plan-Neofluar 10x/0.3 M27 objective on a Zeiss Axiocam 506 mono camera. Raw CZI images were stitched using Zen Blue 2012, and exported as JPEGs for downstream data processing. Tissue sections were permeabilized for 10 minutes to facilitate release of poly-adenylated mRNA and its hybridization to adjacent barcoded capture probes. The optimal permeabilization time for the mouse spinal cord was determined based on permeabilization time course experiments performed using the Visium Tissue Optimization protocol (10X Genomics, CG000238 Rev E). Reverse transcription was performed to generate cDNA incorporating spatial barcodes and unique molecular identifiers (UMIs). Following second strand synthesis, cDNA was collected and amplified. The number of cDNA amplification cycles for each Visium array was determined by qPCR, as instructed by 10X Genomics.

Visium libraries were constructed using unique Illumina-compatible PCR primers with Dual Index Kit TT Set A (10X Genomics, #1000215). The size distributions of the libraries were assessed by running Agilent High Sensitivity DNA chips on an Agilent 2100 Bioanalyzer instrument. Any libraries containing primer-dimer contaminants, or libraries that fell outside the expected size range (450 ± 50 bp) were discarded and reprocessed. Total library yields were quantified on a Qubit 3.0 fluorometer, followed by final qPCR quantification using the KAPA Library Quantification Kit for Illumina Platforms (KAPA Biosystems). Visium libraries were each diluted to the same concentration and pooled at a concentration of 6 nM prior to sequencing. Pooled Visium libraries were loaded and sequenced on the NovaSeq 6000 using the Illumina Novaseq 6000 S4 Reagent v1.5 Kit (200 cycles). The final library pool was sequenced using the following run recipe: read 1: 100 cycles, read 2: 100 cycles, i7 index read: 10 cycles, i5 index read: 10 cycles.

#### 10X Genomics Visium Data Pre-processing

Raw FASTQ files and H&E images were processed using Spaceranger v2.0.0, which uses the modified STAR v2.7.2a pipeline for genome alignment and performs barcode/UMI counting to generate feature-spot matrices. Reads were aligned to the mouse reference genome (GRCm38.p6/mm10) using the GENCODE vM25 gene annotation. Following alignment, we observed an average of 1706 median genes and 3641 Median UMI transcripts per ST spot across our dataset of 38,696 total ST spots. Spaceranger metrics can be found in Table S2.

#### Spinal Cord Histology Annotation

The Visium spots within each tissue section were manually assigned to one of eleven anatomical annotation regions (AARs) using Loupe Browser (10X Genomics, version 8.0), based on H&E-stained gross tissue morphology, cellular composition, and established anatomical landmarks^48^. These annotations enabled comparisons of regional transcriptomic changes between age groups. Tissue was first partitioned into grey matter, identified by its characteristic butterfly-shaped morphology and lighter eosin staining, and surrounding white matter, distinguished by its darker pink/purple staining and lower cellular density. For annotation purposes, the grey matter was subdivided into four regions: 1) the central canal, identified by its centrally located cluster of small, densely packed nuclei; 2) the dorsal horn, containing a heterogeneous population of cells with variable nuclear sizes; 3) the ventral horn, identified by the presence of large lower motor neuron cell bodies; and 4) the medial grey matter, located between the dorsal and ventral horns and surrounding the central canal. The white matter was subdivided into: 5) the dorsal medial white matter, located between the dorsal horns and containing ascending dorsal column fibers; 6) the medial lateral white matter, situated between the dorsal and ventral horns adjacent to the medial grey matter; 7) the ventral medial white matter, located between the ventral horns; 8) the ventral lateral white matter, positioned lateral to the ventral horns; and (9-11) the dorsal, lateral and ventral edge regions, which encompassed the outer borders of the corresponding white matter regions. Spots overlying tissue artifacts, including folds and tears were excluded from downstream analysis. Following annotation, the final dataset contains 38,696 individual spots across 20 arrays.

#### Spatial Data Integration and UMAP Embedding

To account for differences in count depth across spots, the transcript counts within individual spots were normalized by dividing all transcript counts in a spot by a size factor to reach the median count depth across all spots. To mitigate the risk of over-inflating low-count spots, size factors were clipped to a minimum value of 0.25. These region-normalized counts were log1p-transformed and scaled by dividing by the interquartile range (RobustScaler). Genes that had a variance > 1 after scaling were rescaled so that their variance was equal to 1, resulting in standardized spatial counts. PCA was performed on the standardized counts using a truncated singular value decomposition with 25 components. Spots were embedded into a 25-neighbor k-NN graph using these 25 components. The spots were then projected into 2 dimensions with the UMAP algorithm using the k-NN embedding (minimal distance of 0.5; initialized from spectral graph embedding).

#### Splotch Modeling of Spatial Gene Expression Across Age Groups

Raw sequencing data was filtered to remove mitochondrially encoded genes, lncRNAs, pseudogenes, and genes expressed in less than 1% of spatial spots. Any spots containing fewer than 100 UMI transcript counts were filtered out prior to Splotch modeling. In addition, any spots without an assigned annotation label or without at least one immediate neighbor in the Visium grid were discarded to prevent singularities in the spatial autocorrelation component of the Splotch model. The final dataset consists of expression data for 13,280 genes measured across 38,696 spatial spots across all mice, with an average sequencing depth of 3641 UMIs per spot.

The Splotch model enables the robust measurement of expression differences between different age groups and anatomical regions and is designed to rigorously account for the uncertainty of low gene counts. Splotch can analyze multiple tissue sections simultaneously, stratified by age group, enabling quantification of biological and technical variation. Splotch modeling was performed as previously described^46–48,54–57^. Briefly, Splotch applies a zero-inflated Poisson likelihood function (where the zero inflation accounts for technical dropouts), to model the expression (λ) of each gene in each spot. This expression level λ is modeled using a generalized linear model (GLM) with three components: the characteristic expression of a spot’s AAR (β), autocorrelation with the spot’s immediate spatial neighbors (Ψ), and spot-level variation (ε). Moreover, the characteristic gene expression rate (β) is hierarchically formulated to account for the experimental design, with donor-level effects (level 3) conditioned on sex (level 2) and further on age group (level 1), to allow for testing the effect of different sample covariates on differential gene expression. β is measured in log-scaled counts per spot within a given AAR. Performing posterior inference on the model using the observed spatial gene expression data enables the identification of age-associated group level trends in spatial gene expression (β) that best explain our observations.

#### Splotch Spatial Differential Gene Expression Analysis

To measure differences in expression between two sample conditions using Splotch, we examined the posterior expression distributions (β) for each annotated region between pairs of mouse age groups (i.e. 24 months vs 6 months). The Savage-Dickey density ratio of the likelihood of the null hypothesis under the prior vs posterior model was computed to estimate the Bayes factor (BF) for each gene in a given pairwise comparison (for instance, the expression of ‘Lgals3’ in the ventral medial white matter between 24-month vs 6-month-old mice). A threshold of BF ≥ 3 and a mean expression level of β ≥ -5 in at least one condition was used to identify a gene as differentially expressed between two conditions. Bayes factor tables with summary statistics (including BF, log2-fold change, β_age group 1_, β_age group 2_) and a full list of differentially expressed genes in each annotated region and pairwise comparison can be found in Tables S10 and S11, respectively.

#### Identification of Spatial Gene Co-expression Modules and Submodules

Spatial gene co-expression modules were identified as previously described^46,47,54,56^. Briefly, all the mean posterior spot-level gene expression estimates (λ) from Splotch were used to construct a matrix of 38,696 rows (spots) and 13,280 columns (genes). The λ values were capped at the 99^th^ percentile to reduce the influence of outliers and scaled by dividing by the gene interquartile range. Any genes with high variability (variance > 1) were re-scaled to have their variance equal to 1. Next, Pearson correlation coefficients were computed for each gene pair across all spots, resulting in a 13,280 by 13,280 correlation matrix. A correlation distance (1 – Pearson correlation coefficient) was calculated for each gene pair and used as a distance metric for a k-NN graph embedding (k = 10 neighbors). Genes were projected into 2 dimensions using the UMAP algorithm and grouped into modules using Leiden clustering (resolution = 2).

For each spatial module, cell-type associated submodules were identified using orthogonal snRNA-seq data. Raw snRNA-seq count data was depth-normalized to 10,000 counts/nucleus and then log1p-transformed. Within each individual module, Pearson correlations were calculated for each gene pair using the transformed snRNA-seq data. The snRNA-seq correlation distance (1 – Pearson correlation coefficient) for each gene pair was used as the distance metric for k-NN graph embedding (k = 10 neighbors). Genes with similar co-expression patterns across nuclei were clustered into submodules using the Leiden algorithm (resolution = 0.75).

Python functions for spatial module and submodule identification can be found in the scself package (single-cell self supervised; version 0.4.8). Genes within each spatial gene module and submodule can be found in Table S7.

#### Module Scores

A module score, or the combined scaled expression of all genes comprising a module, was calculated for each spot. For each gene, the modeled Splotch λ expression was linearly scaled to a min-max range of 0-1, where 0 is set to the 1^st^ percentile and 1 is set to the 99^th^ percentile. Any value below the 1^st^ percentile was clipped to 0, and any value above the 99^th^ percentile was clipped to 1. For each spatial module, a module score was calculated for each spot by computing the mean of the 0-1 scaled scores for each gene in the module. Spot-level module scores were then aggregated by computing the median module score for all spots in each anatomical annotation region (AAR) for each array and rescaling the median AAR level module scores to a 0-1 range. These scaled AAR-array module scores were used for statistical testing. Welch’s t-test was used to compare module scores across different age groups, correcting for false discovery with Benjamini-Yekutieli (FDR-BY). The module scores for each AAR within each array were further aggregated into mean module scores per age group for plotting.

#### Gene Ontology Enrichment Analysis

Gene ontology (GO) enrichment was performed using g:Profiler, by testing input gene lists for overrepresentation of biological terms relative to a specified background gene set. Enrichment significance was assessed using a hypergeometric test, with p-values adjusted for multiple hypothesis testing. Gene lists consisted of genes assigned to each spatial module, with the background set comprising all genes included in the Splotch spatial model. Gene ontology enrichment terms for the mouse spatial gene modules and submodules can be found in Table S9.

### SINGLE NUCLEUS RNA SEQUENCING (snRNA-seq)

#### Mouse snRNA-seq Data Processing

Published 10X Genomics snRNA-seq sequencing data from five mouse spinal cord studies was acquired from NCBI Gene Expression Omnibus (GEO: GSE161621^49^, GEO: GSE167597^50^, GEO: GSE184370^51^, GEO: GSE172167^52^, GEO: GSE248684^53^). Sequencing data from all five studies were independently processed using a standardized alignment pipeline, with reads aligned to the mouse reference genome (GRCm38.p6/mm10) using the GENCODE vM25 gene annotation, and Cellranger v9.0.0 to generate gene expression count matrices. Cellranger QC metrics can be found in Table S5. Each dataset was subsequently preprocessed independently.

Individual nuclei were retained in each dataset if they had between 500 and 50,000 gene expression transcripts (UMIs), and a maximum fraction of mitochondrially-encoded transcripts of 0.2. To account for differences in sequencing depth across nuclei, raw transcript counts were normalized to a target sum 10,000 counts per nucleus. This was achieved by dividing all the transcript counts for each nucleus by a size factor, to reach a total of 10,000 counts per nucleus. Mitochondrially encoded genes (*Mt*-) and *Malat1* were excluded from the size factor calculations, but were otherwise retained for downstream analysis. Normalized counts were log(x+1) transformed, and each gene was then scaled by dividing by the interquartile range (RobustScaler). Genes having a variance greater than 1 after scaling were further scaled so that their variance was equal to 1, resulting in log-standardized counts.

Principal Component Analysis (PCA) was performed on the log-standardized counts using a truncated singular value decomposition with 25 components. Nuclei were embedded into a 25 neighbor k-Nearest Neighbors (k-NN) graph using these 25 components. Nuclei were then projected into 2 dimensions with Uniform Manifold Approximation and Projection (UMAP) using the k-NN embedding (minimal distance of 0.5; initialized from spectral graph embedding).

#### Mouse Cell Type Annotation and Data Integration

Nuclei from each of the five mouse snRNA-seq datasets were clustered and annotated independently. Following pre-processing, nuclei were clustered into a large number of groups with the leiden algorithm (resolution = 5). Each cluster was assigned to one of 8 broad cell types (excitatory neuron, inhibitory neuron, motor neuron, oligodendrocyte, oligodendrocyte progenitor cell, astrocyte, microglia, endothelial) based on the relative expression of canonical marker genes (Table S6). Clusters exhibiting co-expression of gene markers from multiple broad cell types were labeled as a ‘Doublet’ and excluded from further analysis. Clusters with low overall UMI counts or low marker expression were classified as low-quality nuclei and were also discarded.

Following annotation, the five datasets were concatenated into a single AnnData object. Gene expression counts were subsequently renormalized, log(x+1) transformed, and scaled as described above. PCA was performed on the resulting log-standardized expression matrix, and the first 25 principal components were used to construct a k-NN graph. Nuclei were then projected into 2 dimensions with UMAP.

The final dataset contained 129,564 high-quality nuclei for downstream analysis.

#### Nuclei Isolation from Human Post-mortem Spinal Cord Tissue

Nuclei were isolated from fresh frozen samples using a modified protocol^47,54,56,77,78^. Approximately 150 mg of frozen tissue was collected per sample into pre-chilled 1.5 mL Eppendorf tubes. Samples received 700 µL of homogenization buffer (320 mM sucrose, 5 mM CaCl_2_ [Millipore Sigma; #21115], 3 mM magnesium acetate [Thermo Fisher; AAJ60041AE], 10 mM Tris-HCl pH 7.8 [prepared from Thermo Fisher’s #AM9850G and #AM9855G], 0.1 mM EDTA pH 8 [Fisher; 15-575-020], 0.1% IGEPAL CA-630 [Millipore Sigma, I8896-50ML], 1 mM DTT [Millipore Sigma, 646563-10X], 0.4 U/µL RNAse inhibitor [Millipore Sigma, 03335399001], H_2_O) and incubated on ice for 5 minutes. Samples were homogenized in 1 mL Dounce tissue grinders (DWK Life Sciences, 357538) using 10-15 strokes with the loose pestle followed by 10-15 strokes with the tight pestle. After a 5-minute incubation on ice, homogenized samples were filtered through a 70 µm strainer (Bel-Art, H13680-0070) into 2 mL Eppendorf tubes and mixed with an equal volume of working solution (50% OptiPrep [STEMCELL Technologies, #07820], 5 mM CaCl_2_, 3 mM magnesium acetate, 10 mM Tris-HCl pH 7.8, 0.1 mM EDTA pH 8, 1 mM DTT). For the OptiPrep gradient, 750 µL of 30% OptiPrep solution (30% OptiPrep, 134 mM sucrose, 5 mM CaCl_2_, 3 mM magnesium acetate, 10 mM Tris-HCl pH 7.8, 0.1 mM EDTA pH 8, 0.04% IGEPAL CA-630, 1 mM DTT, 0.17 U/µL RNAse inhibitor) was added over 300 µL 40% OptiPrep solution (40% OptiPrep, 96 mM sucrose, 5 mM CaCl_2_, 3 mM magnesium acetate, 10 mM Tris-HCl pH 7.8, 0.1 mM EDTA pH 8, 0.03% IGEPAL CA-630, 1 mM DTT, 0.12 U/µL RNAse inhibitor) in fresh 15 mL Falcon tubes. Homogenized samples were gently layered onto the gradient and centrifuged at 3,000 RCF for 20 minutes at 4°C in a swinging-bucket rotor with the brake off. Approximately 150 µL of nuclei were collected from the 30%/40% interphase and transferred to new 1.5 mL Eppendorf tubes. Samples received 500 µL of wash buffer (1% BSA, 1 U/µL RNAse inhibitor, PBS) and incubated on ice for 5 minutes. Nuclei were gently resuspended by pipetting and centrifuged at 500 RCF for 5 minutes at 4°C in a fixed-angle rotor. The supernatant was removed, and nuclei were resuspended in ∼10-20 µL of diluted 10x Genomics Nuclei Buffer. For nuclei counting, 1 µL of the nuclei suspension was mixed with 4 µL of PBS and 5 µL of 125 nM SYTOX (Thermo Fisher; #S7020), incubated for 5 minutes, and assessed for GFP^+^ nuclei using the Countess II FL instrument (Thermo Fisher, #AMQAF1000). Each suspension was diluted to 3,230-8,060 nuclei/µL for a target recovery of 10,000 nuclei per sample.

#### 10X Genomics snRNA-seq Library Preparation and Sequencing

Single nucleus RNA-seq and snATAC-seq libraries were generated with the Chromium Next GEM Single Cell Multiome ATAC + Gene Expression Kit (10x Genomics, #PN-1000283) following the Chromium Next GEM Single Cell Multiome ATAC + Gene Expression User Guide (CG000338 Rev F). Quality control of cDNAs and libraries was assessed using a Qubit Flex Fluorometer (Thermo Fisher, Q33327) and an Agilent 2100 Bioanalyzer (Agilent High Sensitivity DNA Kit, 5067-4626). Library quantification was performed using the KAPA Library Quantification Kit for Illumina Platforms (Kapa Biosystems, 07960255001). Single nucleus ATAC-seq libraries were pooled equimolarly and sequenced with the following run configurations: Read 1: 50 cycles; Read 2: 49 cycles; i5: 24 cycles; i7 Index: 8 cycles each. Single nucleus RNA-seq libraries were pooled equimolarly and sequenced with the following run configurations: Read 1: 28 cycles; Read 2: 90 cycles; i5 and i7 Index: 10 cycles each. ATAC-seq data was not analyzed as it was outside the scope of this study.

#### Human snRNA-seq Data Processing

Single nucleus RNA-seq libraries were aligned using the GRCh38 reference genome (hg38; GENCODE v32; Cellranger human reference 2020-A), and a count matrix was generated using Cellranger v9.0.0. Cellranger QC metrics can be found in Table S5. Individual nuclei were retained in the dataset if they had between 500 and 50,000 gene expression transcripts (UMIs), and a maximum fraction of mitochondrially-encoded transcripts of 0.2. A total of 281,726 nuclei from 25 libraries passed this initial filtering.

To account for differences in sequencing depth across nuclei, raw transcript counts were normalized to a target sum 10,000 counts per nucleus. This was achieved by dividing all the transcript counts for each nucleus by a size factor, to reach a total of 10,000 counts per nucleus. Mitochondrially encoded genes (MT-) and *MALAT1* were excluded from the size factor calculations, but were otherwise retained for downstream analysis. Normalized counts were log(x+1) transformed, and each gene was then scaled by dividing by the interquartile range (RobustScaler). Genes having a variance greater than 1 after scaling were further scaled so that their variance was equal to 1, resulting in log-standardized counts.

Principal Component Analysis (PCA) was performed on the log-standardized counts using a truncated singular value decomposition with 25 components. Nuclei were embedded into a 25 neighbor k-Nearest Neighbors (k-NN) graph using these 25 components. Nuclei were then projected into 2 dimensions with Uniform Manifold Approximation and Projection (UMAP) using the k-NN embedding (minimal distance of 0.5; initialized from spectral graph embedding).

#### Human Cell Type Annotation

All pre-processed nuclei were clustered into a large number of groups with the leiden algorithm (resolution = 5). Each cluster was assigned to one of 10 broad cell types (excitatory neuron, inhibitory neuron, motor neuron, serotonergic neuron, oligodendrocyte, oligodendrocyte progenitor cell, astrocyte, microglia/immune, endothelial, and meninges) based on the relative expression of canonical marker genes. Annotation gene markers can be found in Table S4. Any cluster co-expressing genes from multiple broad cell types was labeled as a ‘Doublet’ and removed. Clusters with low overall UMI counts or low marker expression were classified as low-quality nuclei and were also discarded.

Motor neurons were subjected to further quality control following the identification of a substantial population of putative “nuclei” exhibiting high mitochondrially encoded transcript fractions and transcriptional profiles resembling motor neurons, but lacking nuclear-enriched transcript makers (e.g. *MALAT1*), suggesting that these represented cellular debris rather than intact nuclei. Annotated motor neurons were subclustered (resolution = 3) and the expression of mitochondrially encoded (MT-) transcripts and *MALAT1* was assessed within each cluster. Only clusters with low MT-expression but high *MALAT1* expression were retained.

The final dataset contained 198,191 high-quality nuclei for downstream analysis.

#### Cell Proportions Per Sample

Cell type proportions were calculated for each donor by dividing the number of nuclei assigned to a given cell type by the total number of nuclei for that donor. These proportions were then averaged across donors within each age group. Differences in cell-type proportions between age groups were assessed using Welch’s t-test, with Benjamini-Hochberg correction for multiple comparisons.

#### Cell Type-Specific Analysis of Age-Associated Gene Expression

To assess age-associated changes in gene expression, transcript counts were aggregated across nuclei of the same cell type within each donor to generate cell-type specific pseudobulk expression profiles. Gene expression was normalized to transcripts per million (TPM), and expression levels were plotted against donor age for each cell type. Pearson correlation analysis was performed to assess the relationship between gene expression and donor age, with correlation coefficients (r), coefficients of determination (R^2^), and associated p-values were calculated for each comparison. P-values were adjusted for multiple testing using the Benjamini-Hochberg false discovery rate (FDR) correction.

### SERIAL IMMUNOFLUORESCENCE IMAGING USING LUNAPHORE COMET^TM^ (COMET)

#### Tissue Preparation (Sectioning & Fixation)

Mouse and human lumbar spinal cord tissue blocks were embedded in Tissue Plus OCT Compound (Fisher Healthcare; #4585) and sectioned in a cryostat at -16°C. 10 um-thick sections were collected onto Superfrost Plus slides from fresh frozen mouse and human lumbar spinal cords. For mouse imaging, four mice (2 males, 2 females) per age group (6, 12, 18, 24, 30 months) were included, and two mice with two replicate sections each were sectioned onto one slide. For human imaging, 13 donors were included, and one tissue section per slide was collected for each donor (Table S1).

Mouse tissues were fixed prior to COMET imaging following Lunaphore’s recommended preprocessing steps. Slides were thawed at room temperature for 20 minutes, then incubated in Formaldehyde 4% for 40 minutes on an orbital shaker. Slides were then washed for 5 minutes at room temperature in Lunaphore’s Multistaining Buffer (MSB; Advanced Cell Diagnostics; #BU09) on a shaker. Finally, slides were incubated for 20 minutes at room temperature in Tris-Buffered Saline (TBS) with 0.2% Triton on a shaker and placed in MSB until loading into the COMET machine.

Slides containing human tissue were thawed at room temperature for 7 minutes, incubated in 10% Neutral Buffered Formalin for 40 minutes, and washed 3 times in MSB (5 min/wash on a shaker). Slides were then incubated in Tris-Buffered Saline (TBS) with 0.1% Triton for 12.5 minutes on a shaker followed by Tris-Buffered Saline (TBS) with 0.1% Tween for 5 minutes on a shaker and placed in MSB until loading into the COMET machine.

#### COMET Multi-plex Immunofluorescence Imaging

The COMET^TM^ platform (Lunaphore Technologies) was used for sequential immunofluorescence staining according to a COMET software-generated protocol. Staining, imaging, elution, quenching, and wash conditions were performed as shown in Table S13. For mouse, all primary and secondary antibodies were diluted in MSB. For human, all primary and secondary antibodies were diluted in Intercept Antibody Diluent (LICORbio; 927-65001). All primary antibodies were incubated for 4 minutes at 27°, with the exception of p21 and p16 in human (16 minutes), and all secondary antibodies were incubated 2 minutes at 27°, and elution at 37°. To assess for autofluorescence, non-specific secondary antibody signal, and residual signal following elution, secondary-only control cycles and post-elution images were included.

#### Image Preprocessing and Export

The COMET software automatically aligned and flat-field corrected images, and image stacks were exported as 12-bit OME-TIFF files. Horizon Viewer was used to open images, and secondary antibody control images were used for background correction. Images were then exported for downstream analysis.

#### Anatomical Layer Annotation

Images were imported into HALO v4.4.7276.136, then annotated using the annotation tool. Spinal cord subregions were defined using DAPI, neuronal cell markers (NFH and SMI-32), and a white matter marker (MBP). White (dorsal medial white matter, medial lateral white matter, ventral medial white matter, ventral lateral white matter) vs. grey matter (dorsal horn, medial grey matter, ventral horn) regions were designated based on the border between the neuronal cell markers and MBP. Grey and white matter regions were further subdivided as described above, excluding edge regions. Areas with overlying tissue artifacts, including folds and tears, or that were out of focus during imaging were excluded from downstream analysis.

#### Image Analysis in HALO^®^

For area quantification analysis in Figure S3, the Area Quantification module was used to quantify area positive for APOE staining. A single intensity threshold for positive staining was set and applied to all images, regions, and animals. The threshold was chosen based on pixel-intensity histograms and a threshold sweep. Results were unchanged between thresholds at 800-1000 (Table S13).

For IBA1 and P2Y12 area analysis in Figure 5, the Object Colocalization module was used to detect percent area coverage for IBA1, P2Y12, and IBA+P2Y12 co-localization. To restrict analysis to parenchymal microglia and avoid contribution from vessel-associated signal, areas that were double positive for vascular markers (Claudin-5, Collagen IV, CD31, and aSMA) and IBA1 or P2Y12 were excluded from analysis. A single intensity threshold was used for all markers.

The Highplex FL module was used to quantify microglia. HALO’s pretrained nuclei segmentation model was used to detect nuclei, and an in-house membrane segmentation model trained through HALO AI was used to improve cell-boundary detection for microglia^47^. Microglia were defined as IBA1+, P2Y12+, or IBA1 and P2Y12+, and all microglia had to be DAPI+ and negative for NeuN, OLIG2, Claudin-5, Collagen IV, CD31, and aSMA. Welch’s t-test with Benjamini-Hochberg correction was used for pairwise comparisons between age groups.

### GENERAL SOFTWARE AND PROCEDURES

Python, scanpy, and matplotlib were used to generate all figures and plots. Figures were constructed using Adobe Illustrator, and Figure 1A and 2A were designed using Biorender.

### Data and Code Availability

Raw sequencing (Visium, Multiome) and imaging (COMET) data will be available through the SenNet Consortium data portal upon publication: https://doi.org/10.60586/SNT655.SBBT.369.

Code used to analyze and visualize data is available upon request.

## Acknowledgements

This work was supported by awards from the National Institutes of Health (NIH) (RO1NS116350 and RO1NS118183), the National Institute of Aging (NIA) (U54AG076040 (PI:Phatnani) and AG075932 (PI: Schilling)) and TargetALS (FD-2023-GEN-S1).

Computational resources were provided by the New York Genome Center. We thank Mariecia Pook for helping with project coordination, Dr. Neil A. Shneider for providing us with the mice used in our spatial transcriptomics experiments, and the San Diego Tissue Mapping Center and Dr. Marissa Shaefer’s Lab at the Mayo Clinic for generously sharing their curated senescence gene lists. Finally, we would like to acknowledge all the donors who contributed the tissues that were used in this study.

## Author Contributions

Conceptualization-BS, HP

Methodology-OK, MX, ACD, CAJ, IH, NB, MAW, CDK, MB, PS

Software-ACD, CAJ

Investigation-OK, MX

Formal analysis-OK, MX, ACD, CAJ

Data upload-CM, MHP

Writing – original draft preparation-OK, MX

Writing – review & editing-All authors

Resources-NAS, BS, HP

Supervision-BS, HP

Funding acquisition-VM, BS, HP

