## Supplemental Figures 1-7 for "Spatiotemporal profiling reveals emergence of cell type-specific molecular aging and senescence programs in the spinal cord"

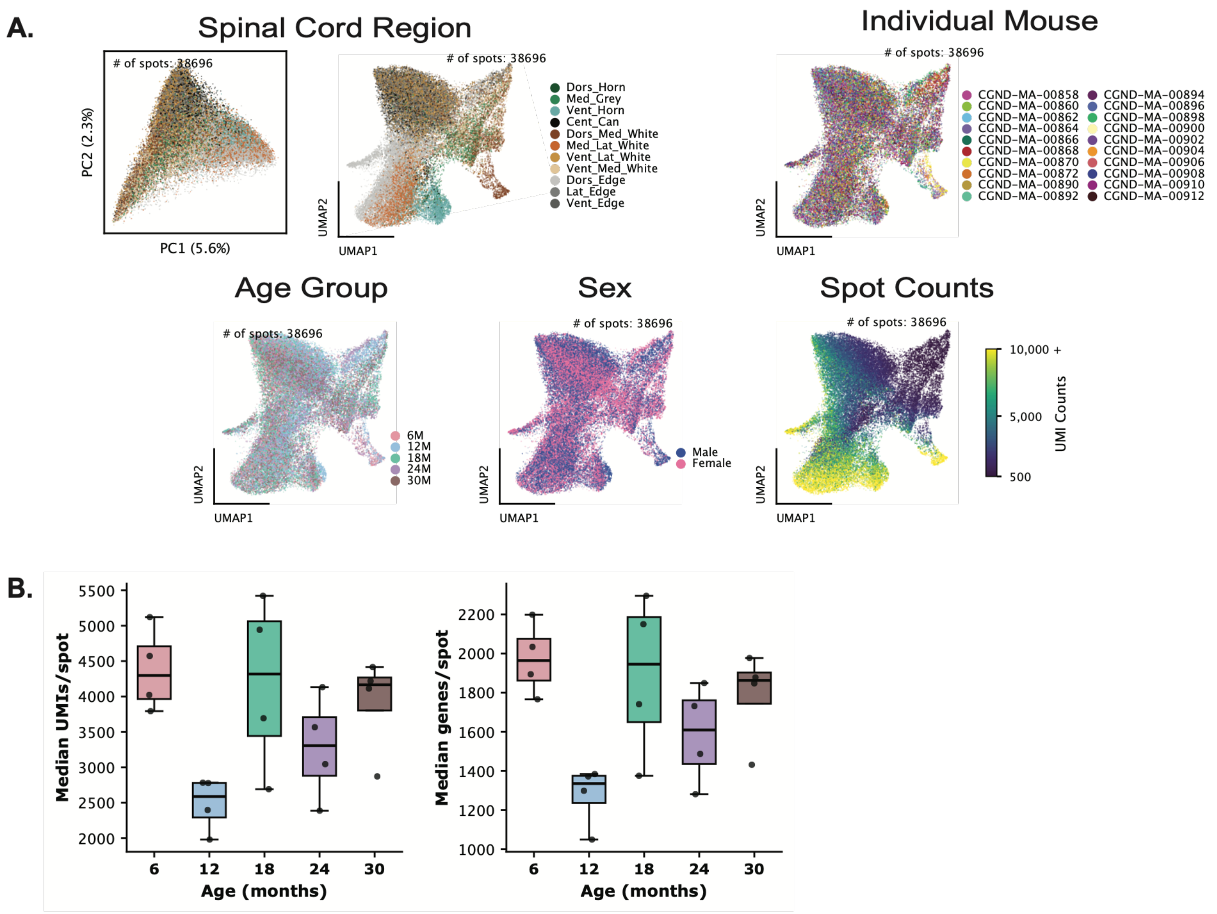
A.
Spinal Cord Region
### of spots: 38696
PC1 (5.6%)
PC2 (2.3%)
UMAP1
UMAP2
Dors_Horn
Med_Grey
Vent_Horn
Cent_Can
Dors_Med_White
Med_Lat_White
Vent_Lat_White
Vent_Med_White
Dors_Edge
Lat_Edge
Vent_Edge
Individual Mouse
CGND-MA-00858
CGND-MA-00860
CGND-MA-00862
CGND-MA-00864
CGND-MA-00866
CGND-MA-00868
CGND-MA-00870
CGND-MA-00872
CGND-MA-00890
CGND-MA-00892
CGND-MA-00894
CGND-MA-00896
CGND-MA-00898
CGND-MA-00900
CGND-MA-00902
CGND-MA-00904
CGND-MA-00906
CGND-MA-00908
CGND-MA-00910
CGND-MA-00912
Age Group
6M
12M
18M
24M
30M
Sex
Male
Female
Spot Counts
UMI Counts
10,000 +
5,000
500
B.
Median UMIs/spot
Median genes/spot
Age (months)
6
12
18
24
30

**Figure S1. Mouse Visium ST sequencing quality.**

**A)** PCA and UMAP plots depict all spots colored by spinal cord region annotation, individual mouse, age group, sex, and RNA (UMI) counts.

**B)** Differences in QC metrics (Median UMIs/spot, Median genes/spot) between age groups (6, 12, 18, 24, 30 months). Each dot is an individual mouse. No significant differences between age groups across either metric (Mann-Whitney U test).

A.

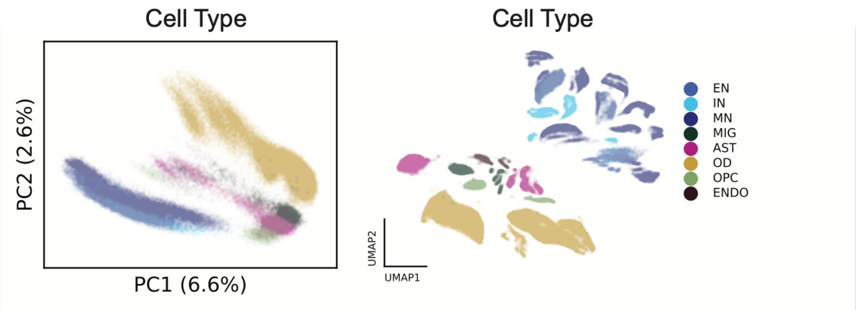


B.

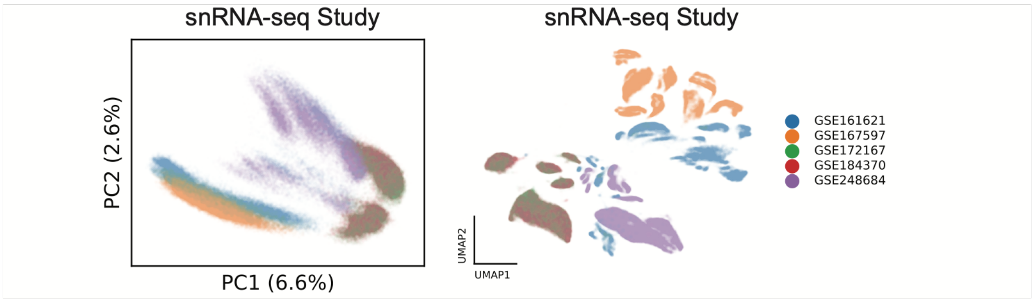


C.

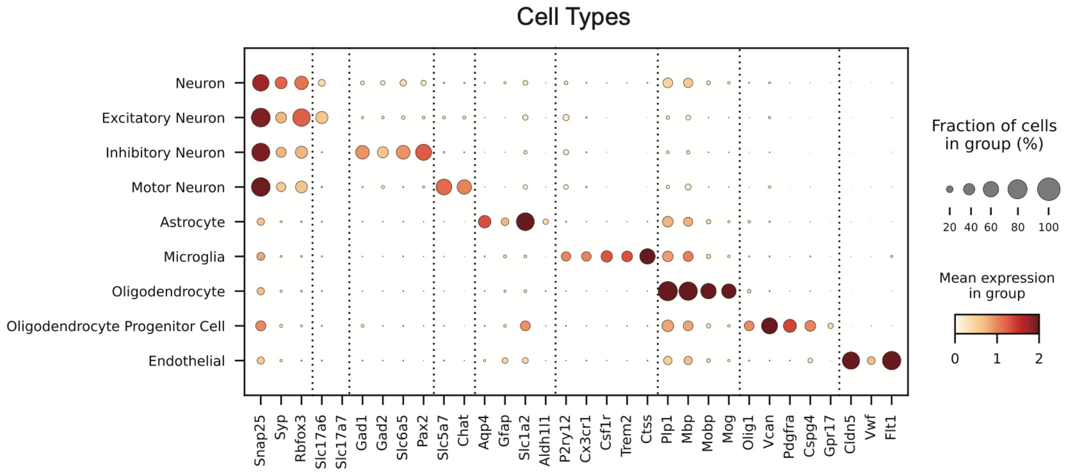


D.

#### Cell Type Counts per Study

| | GSE161621 Blum et al. | GSE167597 Alkaslasi et al. | GSE184370 Kathe et al. | GSE172167 Matson et al. | GSE248684 Iida et al. | Combined |
|---|---|---|---|---|---|---|
| **OD** | 2,230 (9.6%) | 102 (0.4%) | 18,910 (68.4%) | 15,042 (69.7%) | 25,532 (79.0%) | 61,816 (47.7%) |
| **MN** | 12,161 (52.6%) | 11,649 (46.8%) | 0 (0.0%) | 0 (0.0%) | 0 (0.0%) | 23,810 (18.4%) |
| **AST** | 1,110 (4.8%) | 0 (0.0%) | 4,091 (14.8%) | 3,099 (14.4%) | 3,972 (12.3%) | 12,272 (9.5%) |
| **EN** | 3,919 (16.9%) | 8,268 (33.3%) | 0 (0.0%) | 0 (0.0%) | 0 (0.0%) | 12,187 (9.4%) |
| **IN** | 2,661 (11.5%) | 4,846 (19.5%) | 0 (0.0%) | 0 (0.0%) | 0 (0.0%) | 7,507 (5.8%) |
| **MIG** | 212 (0.9%) | 0 (0.0%) | 2,134 (7.7%) | 1,470 (6.8%) | 1,451 (4.5%) | 5,267 (4.1%) |
| **OPC** | 231 (1.0%) | 0 (0.0%) | 1,911 (6.9%) | 1,440 (6.7%) | 1,362 (4.2%) | 4,944 (3.8%) |
| **ENDO** | 606 (2.6%) | 0 (0.0%) | 618 (2.2%) | 537 (2.5%) | 0 (0.0%) | 1,761 (1.4%) |
| **Total** | **23,130** | **24,865** | **27,664** | **21,588** | **32,317** | **129,564** |

**Figure S2. Mouse snRNA-seq reference dataset cell type annotations.**

**A)** PCA and UMAP depicting all nuclei colored by cell type annotation from five mouse spinal cord snRNA-seq studies.

**B)** PCA and UMAP depicting all nuclei colored by their respective mouse spinal cord snRNA-seq study.

**C)** Cell type annotation for nuclei in mouse spinal cord snRNA-seq dataset.

**D)** Cell type counts and proportions per snRNA-seq study.

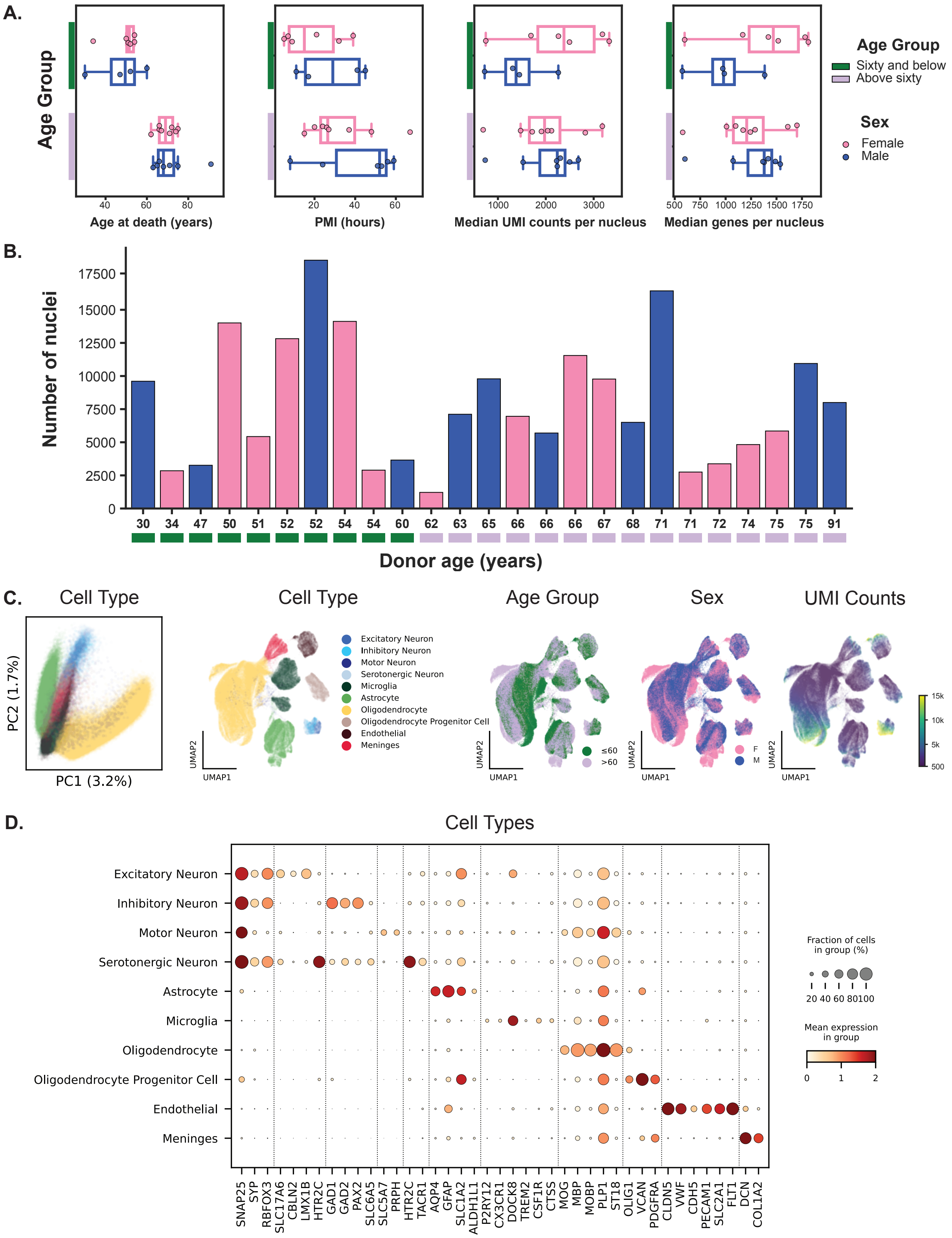
A.
Age Group
Age at death (years)
PMI (hours)
Median UMI counts per nucleus
Median genes per nucleus
Age Group
Sixty and below
Above sixty
Sex
Female
Male
B.
Number of nuclei
Donor age (years)
30 34 47 50 51 52 52 54 54 60 62 63 65 66 66 66 67 68 71 71 72 74 75 75 91
C.
Cell Type
PC2 (1.7%)
PC1 (3.2%)
Cell Type
Excitatory Neuron
Inhibitory Neuron
Motor Neuron
Serotonergic Neuron
Microglia
Astrocyte
Oligodendrocyte
Oligodendrocyte Progenitor Cell
Endothelial
Meninges
Age Group
≤60
>60
Sex
F
M
UMI Counts
UMAP1
UMAP2
D.
Cell Types
Excitatory Neuron
Inhibitory Neuron
Motor Neuron
Serotonergic Neuron
Astrocyte
Microglia
Oligodendrocyte
Oligodendrocyte Progenitor Cell
Endothelial
Meninges
SNAP25 SYP RBFOX3 SLC17A6 CBLN2 LMX1B HTR2C GAD1 GAD2 PAX2 SLC6A5 SLC5A7 PRPH HTR2C TACR1 AQP4 GFAP SLC1A2 ALDH1L1 P2RY12 CX3CR1 DOCK8 TREM2 CSF1R CTSS MOG MBP MOBP PLP1 ST18 OLIG1 VCAN PDGFRA CLDN5 VWF CDH5 PECAM1 SLC2A1 FLT1 DCN COL1A2
Fraction of cells in group (%)
20 40 60 80100
Mean expression in group
0 1 2

**Figure S3. Human snRNA-seq quality and cell type annotation.**

**A)** Differences in metrics between age group or between sexes within each age group plotted against age of death, PMI (hours), and snRNA-seq QC metrics (Median UMI counts/nucleus, Median genes/nucleus). Each dot represents an individual donor. No significant differences between age groups across either metric (Two-sided Mann-Whitney U test, FDR-BH Correction).

**B)** Bar plot depicts total number of nuclei per donor, color coded by age group and sex.

**C)** PCA and UMAP plots depict all nuclei colored by cell type annotation, age group, sex, and RNA (UMI) counts.

**D)** Cell type annotation for nuclei in human spinal cord snRNA-seq data.

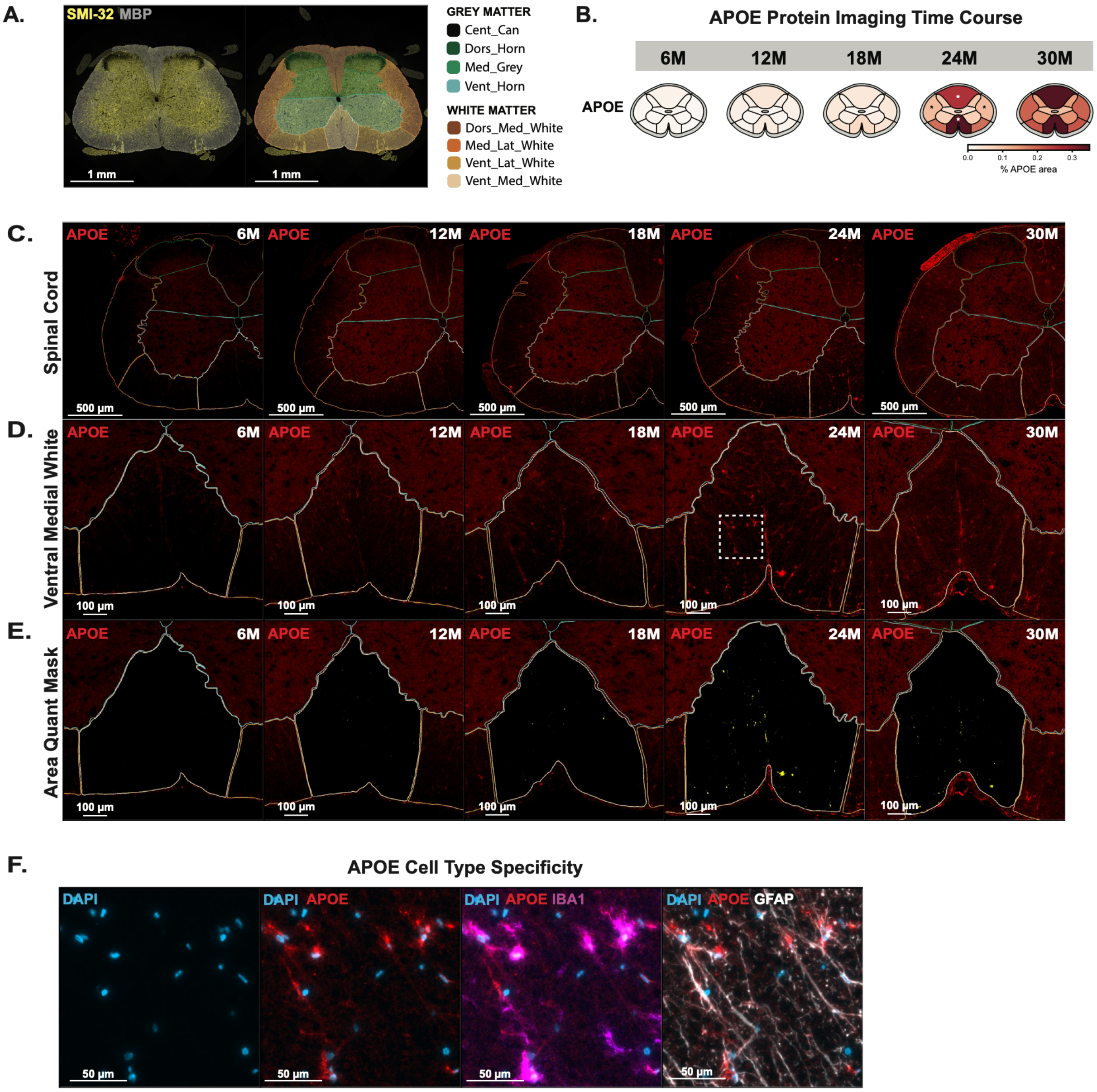
A.
SMI-32 MBP
1 mm
1 mm
GREY MATTER
Cent_Can
Dors_Horn
Med_Grey
Vent_Horn
WHITE MATTER
Dors_Med_White
Med_Lat_White
Vent_Lat_White
Vent_Med_White
B.
APOE Protein Imaging Time Course
6M
12M
18M
24M
30M
APOE
% APOE area
C.
Spinal Cord
APOE
500 µm
D.
Ventral Medial White
100 µm
E.
Area Quant Mask
F.
APOE Cell Type Specificity
DAPI
DAPI APOE
DAPI APOE IBA1
DAPI APOE GFAP
50 µm

**Figure S4. Mouse APOE protein imaging time course.**

**A)** Example mouse spinal cord with spinal cord region annotations overlaid.

**B)** Mean area of APOE in IF as a percentage of total area across each subregion over time course. Asterisk indicates significant differences at time point compared to 6 months (Welch's t-test, Benjamini-Hochberg correction $p < 0.05$).

**C)** Representative IF image of APOE staining at 6, 12, 18, 24, 30 months across half of the spinal cord.

**D)** Representative IF image of APOE staining at 6, 12, 18, 24, 30 months zoomed into the ventral medial white of the spinal cord.

**E)** Area quantification mask for APOE+ staining in the ventral medial white.

**F)** Zoomed in (Box in D) APOE staining overlaid with DAPI, IBA1, and GFAP.

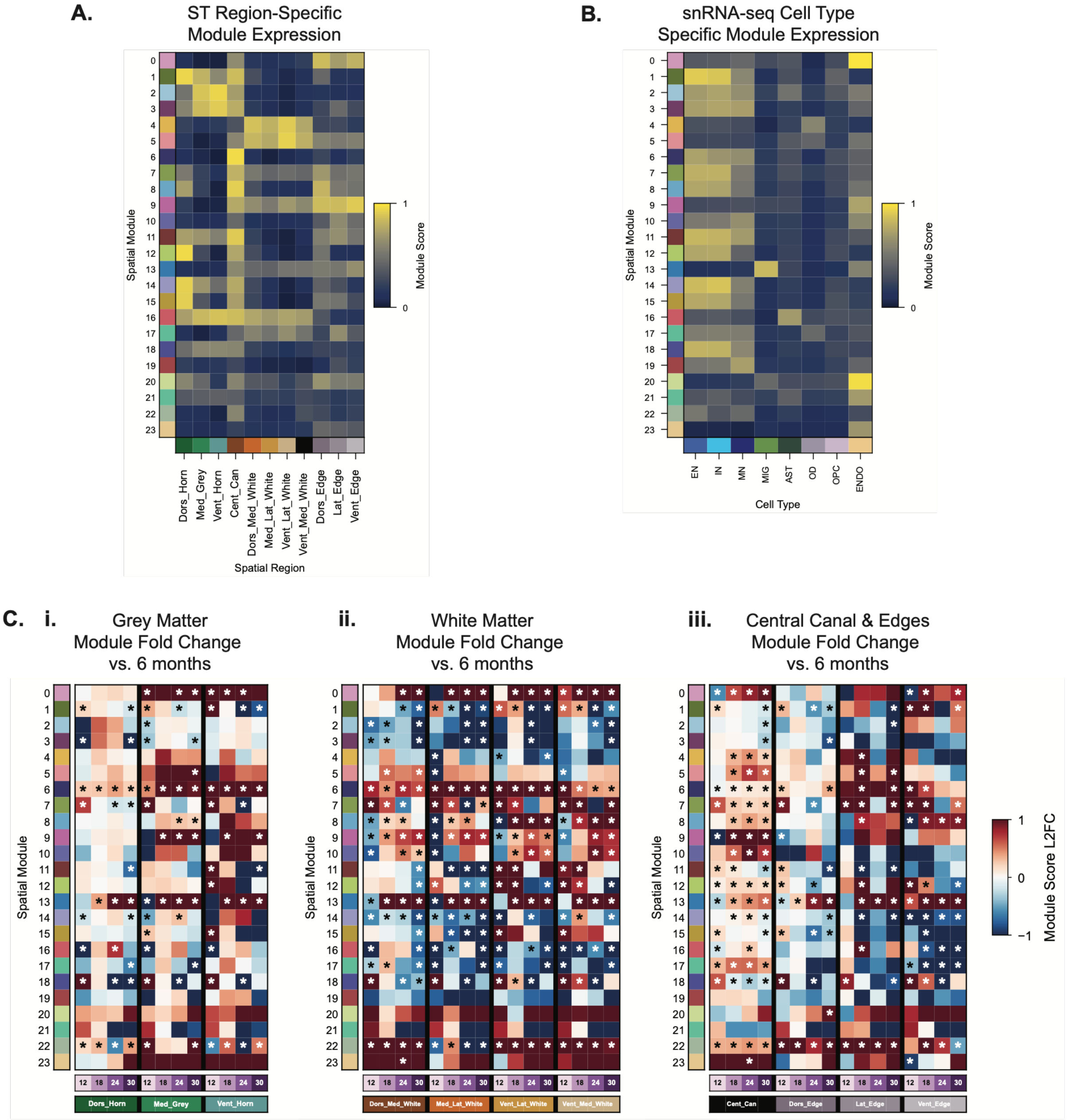

A.
ST Region-Specific Module Expression
Spatial Module
Module Score
Dors_Horn
Med_Grey
Vent_Horn
Cent_Can
Dors_Med_White
Med_Lat_White
Vent_Lat_White
Vent_Med_White
Dors_Edge
Lat_Edge
Vent_Edge
Spatial Region
B.
snRNA-seq Cell Type Specific Module Expression
EN
IN
MN
MIG
AST
OD
OPC
ENDO
Cell Type
C. i.
Grey Matter Module Fold Change vs. 6 months
ii.
White Matter Module Fold Change vs. 6 months
iii.
Central Canal & Edges Module Fold Change vs. 6 months
Module Score L2FC

**Figure S5. Aging trajectories of mouse spatial modules are region- and cell type-specific.**

**A)** Heatmap of module scores across anatomical regions.

**B)** Heatmap of cell type scores for each module in snRNA-seq data.

**C)** Heatmap showing log2 fold change in module scores between 12, 18, 24, 30 month mice vs. 6 month mice in i) grey matter subregions, ii) white matter subregions, and iii) central canal and edge regions. Color indicates log2 fold change of each module at 12-, 18-, 24-, and 30-month time points relative to 6 months, and asterisk indicates significant difference between time points and 6 months (Welch's t-test, Benjamini-Yekutieli correction $p < 0.05$).

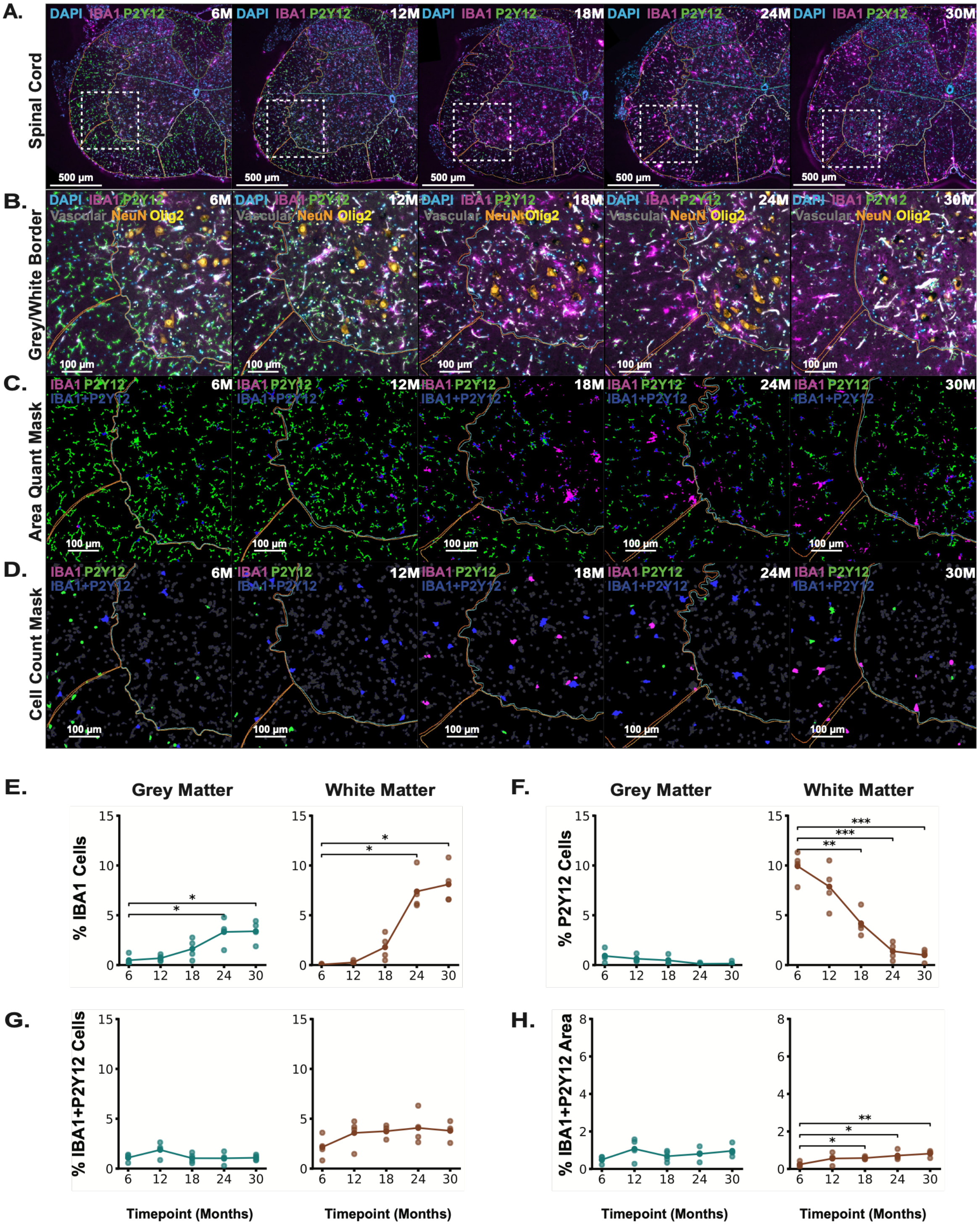
A.
B.
C.
D.
Spinal Cord
Grey/White Border
Area Quant Mask
Cell Count Mask
DAPI IBA1 P2Y12
Vascular NeuN Olig2
IBA1 P2Y12
IBA1+P2Y12
6M
12M
18M
24M
30M
500 µm
100 µm
E.
F.
G.
H.
Grey Matter
White Matter
% IBA1 Cells
% P2Y12 Cells
% IBA1+P2Y12 Cells
% IBA1+P2Y12 Area
Timepoint (Months)

**Figure S6. Mouse IBA1 and P2Y12 area and cell counts with aging.**

**A)** Representative IF image of DAPI, IBA1, and P2Y12 at 6, 12, 18, 24, 30 months across half of the spinal cord.

**B)** Zoomed in (Boxes in A) staining of IBA1 and P2Y12 overlaid with DAPI, Vascular markers (CD31, Collagen IV, Claudin-5, and aSMA), NeuN, and Olig2.

**C)** Area quantification mask for IBA1+, P2Y12+, and IBA1+P2Y12+ staining.

**D)** Cell count mask for IBA1+, P2Y12+, and IBA1+P2Y12+ cells.

**E)** Mean IBA1 cells as a percentage of total cells in grey matter (left) and white matter (right) (Welch's t-test, Benjamini-Hochberg correction $p<0.05$).

**F)** Mean P2Y12 cells as a percentage of total cells in grey matter (left) and white matter (right) (Welch's t-test, Benjamini-Hochberg correction $p<0.05$).

**G)** Mean IBA1+P2Y12 cells as a percentage of total cells in grey matter (left) and white matter (right) (Welch's t-test, Benjamini-Hochberg correction $p<0.05$).

**H)** Mean IBA1+P2Y12 area as a percentage of total area in grey matter (left) and white matter (right) (Welch's t-test, Benjamini-Hochberg correction $p<0.05$).

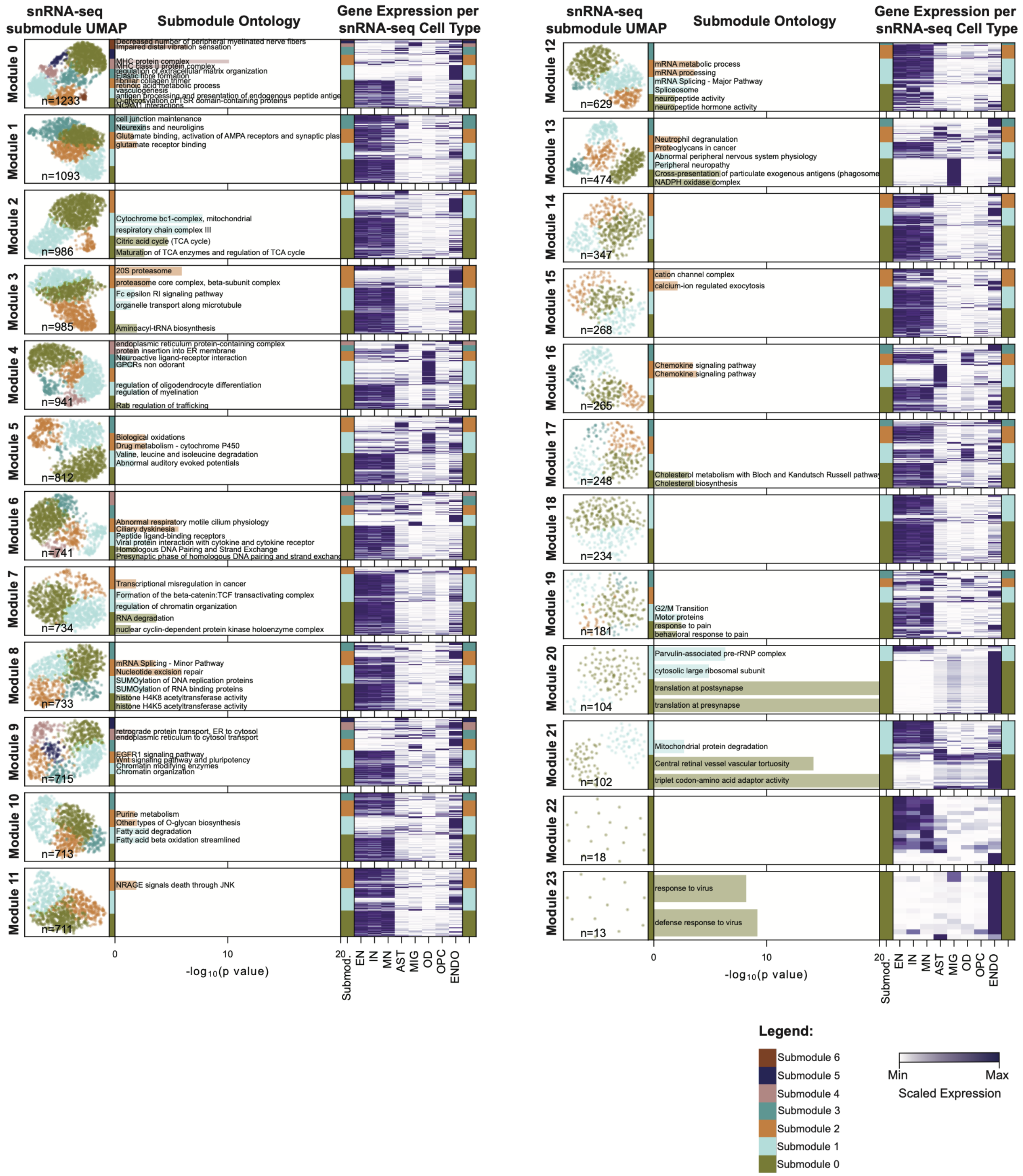

snRNA-seq submodule UMAP
Submodule Ontology
Gene Expression per snRNA-seq Cell Type
Module 0
n=1233
Decreased number of peripheral myelinated nerve fibers
Impaired distal vibration sensation
MHC protein complex
MHC class II protein complex
regulation of extracellular matrix organization
Elastic fibre formation
fibrillar collagen trimer
retinoic acid metabolic process
vasculogenesis
antigen processing and presentation of endogenous peptide antigen
O-glycosylation of TSR domain-containing proteins
NCAM1 interactions
Module 1
n=1093
cell junction maintenance
Neurexins and neuroligins
Glutamate binding, activation of AMPA receptors and synaptic plasticity
glutamate receptor binding
Module 2
n=986
Cytochrome bc1-complex, mitochondrial
respiratory chain complex III
Citric acid cycle (TCA cycle)
Maturation of TCA enzymes and regulation of TCA cycle
Module 3
n=985
20S proteasome
proteasome core complex, beta-subunit complex
Fc epsilon RI signaling pathway
organelle transport along microtubule
Aminoacyl-tRNA biosynthesis
Module 4
n=941
endoplasmic reticulum protein-containing complex
protein insertion into ER membrane
Neuroactive ligand-receptor interaction
GPCRs non odorant
regulation of oligodendrocyte differentiation
regulation of myelination
Rab regulation of trafficking
Module 5
n=812
Biological oxidations
Drug metabolism - cytochrome P450
Valine, leucine and isoleucine degradation
Abnormal auditory evoked potentials
Module 6
n=741
Abnormal respiratory motile cilium physiology
Ciliary dyskinesia
Peptide ligand-binding receptors
Viral protein interaction with cytokine and cytokine receptor
Homologous DNA Pairing and Strand Exchange
Presynaptic phase of homologous DNA pairing and strand exchange
Module 7
n=734
Transcriptional misregulation in cancer
Formation of the beta-catenin:TCF transactivating complex
regulation of chromatin organization
RNA degradation
nuclear cyclin-dependent protein kinase holoenzyme complex
Module 8
n=733
mRNA Splicing - Minor Pathway
Nucleotide excision repair
SUMOylation of DNA replication proteins
SUMOylation of RNA binding proteins
histone H4K8 acetyltransferase activity
histone H4K5 acetyltransferase activity
Module 9
n=715
retrograde protein transport, ER to cytosol
endoplasmic reticulum to cytosol transport
EGFR1 signaling pathway
Wnt signaling pathway and pluripotency
Chromatin modifying enzymes
Chromatin organization
Module 10
n=713
Purine metabolism
Other types of O-glycan biosynthesis
Fatty acid degradation
Fatty acid beta oxidation streamlined
Module 11
n=711
NRAGE signals death through JNK
0
10
20
-log10(p value)
Submod
EN
IN
MN
AST
MIG
OD
OPC
ENDO
Module 12
n=629
mRNA metabolic process
mRNA processing
mRNA Splicing - Major Pathway
Spliceosome
neuropeptide activity
neuropeptide hormone activity
Module 13
n=474
Neutrophil degranulation
Proteoglycans in cancer
Abnormal peripheral nervous system physiology
Peripheral neuropathy
Cross-presentation of particulate exogenous antigens (phagosomes)
NADPH oxidase complex
Module 14
n=347
Module 15
n=268
cation channel complex
calcium-ion regulated exocytosis
Module 16
n=265
Chemokine signaling pathway
Chemokine signaling pathway
Module 17
n=248
Cholesterol metabolism with Bloch and Kandutsch Russell pathways
Cholesterol biosynthesis
Module 18
n=234
Module 19
n=181
G2/M Transition
Motor proteins
response to pain
behavioral response to pain
Module 20
n=104
Parvulin-associated pre-rRNP complex
cytosolic large ribosomal subunit
translation at postsynapse
translation at presynapse
Module 21
n=102
Mitochondrial protein degradation
Central retinal vessel vascular tortuosity
triplet codon-amino acid adaptor activity
Module 22
n=18
Module 23
n=13
response to virus
defense response to virus
Legend:
Submodule 6
Submodule 5
Submodule 4
Submodule 3
Submodule 2
Submodule 1
Submodule 0
Min
Max
Scaled Expression

**Figure S7. Cell type submodules of spatial gene modules using correlation with snRNA-seq.**

For each module, plots display from left to right:

1) Module number
2) Gene UMAP plot, where each dot represents a gene colored by its assigned submodule.
3) Top two GO terms enriched in each submodule. Bar is colored by its corresponding submodule, and its length indicates the -log10(p-value).
4) Heatmap showing expression of module's genes per snRNA-seq cell type, min-max per gene.
